# Single-cell analysis of shared signatures and transcriptional diversity during zebrafish development

**DOI:** 10.1101/2023.03.20.533545

**Authors:** Abhinav Sur, Yiqun Wang, Paulina Capar, Gennady Margolin, Jeffrey A. Farrell

## Abstract

During development, animals generate distinct cell populations with specific identities, functions, and morphologies. We mapped transcriptionally distinct populations across 489,686 cells from 62 stages during wild-type zebrafish embryogenesis and early larval development (3–120 hours post-fertilization). Using these data, we identified the limited catalog of gene expression programs reused across multiple tissues and their cell-type-specific adaptations. We also determined the duration each transcriptional state is present during development and suggest new long-term cycling populations. Focused analyses of non-skeletal muscle and the endoderm identified transcriptional profiles of understudied cell types and subpopulations, including the pneumatic duct, individual intestinal smooth muscle layers, spatially distinct pericyte subpopulations, and homologs of recently discovered human *best4*+ enterocytes. The transcriptional regulators of these populations remain unknown, so we reconstructed gene expression trajectories to suggest candidates. To enable additional discoveries, we make this comprehensive transcriptional atlas of early zebrafish development available through our website, Daniocell.

## Introduction

Animals consist of a collection of cells with beautifully diverse shapes, structures, and functions, and this diversity is rebuilt from scratch during the development of each embryo. One central quest of developmental biology is to understand how this morphological and functional diversity observed in distinct cells relates to transcriptional diversity—how much do transcriptional differences result in distinct morphologies and functions and through what mechanisms? In a crucial step toward addressing this question, single-cell RNAseq (scRNAseq) approaches provide an opportunity to map the transcriptionally distinct populations of cells during development in an unbiased way (Briggs et al. 2018; Farrell et al. 2018; Siebert et al. 2019; Cao et al. 2019; Fincher et al. 2018; Hu et al. 2020; Musser et al. 2021; Plass et al. 2018; Wagner et al. 2018; Lindeboom, Regev, and Teichmann 2021; Rozenblatt-Rosen et al. 2021; Li et al. 2022; Tabula Sapiens et al. 2022). As has been demonstrated repeatedly, scRNAseq-derived molecular cell type catalogs even in well-studied systems often reveal cell types or cell states that have been invisible in previous studies (Farrell et al. 2018; Parikh et al. 2019; Satija et al. 2015; Li, Li, et al. 2016; Shekhar et al. 2016). Additionally, the growing collection of molecular catalogs from diverse animals has enabled systematic and detailed comparisons of cell type transcriptional similarity across species, which mark the first steps toward understanding how transcriptional programs change and new cell types emerge during evolution (Shafer, Sawh, and Schier 2022; Tarashansky et al. 2021; Arendt et al. 2019). Finally, single-cell transcriptomes collected in high-resolution timecourses allow for inference of the temporal changes in gene expression that occur as cells are specified and differentiate, using pseudotime trajectory and related approaches (Qiu et al. 2017; Trapnell et al. 2014; Fan et al. 2016; Wolf et al. 2019; Farrell et al. 2018). These analyses can identify the cascades of gene expression that accompany functional and morphological changes within differentiating cells and are valuable for identifying candidate cell-type or gene expression program regulators to test with reverse genetic approaches. Altogether, single-cell transcriptomic approaches provide a useful complement to classical genetic and embryological study of developmental biology by revealing unexpected complexity, characterizing cells at a whole-transcriptome level instead of with selected marker genes, and helping generate and sharpen hypotheses.

Zebrafish are a powerful model for studying vertebrate embryogenesis, since their genetic tractability, optical clarity, external fertilization, and high fecundity have greatly facilitated developmental screens (Driever et al. 1996; ‘The Zebrafish Issue’ 1996; Nusslein-Volhard 2012; Mullins et al. 2021), rigorous lineage tracing (Carney and Mosimann 2018; Kimmel, Warga, and Schilling 1990; Ho and Kimmel 1993; Helde et al. 1994), and detailed mechanistic studies. Moreover, the high degree of developmental conservation among vertebrates has made zebrafish appealing for modeling many human diseases (Lieschke and Currie 2007; Rubinstein 2003). Here, we present a global view of cell type heterogeneity and transcriptional progression during zebrafish development to complement data acquired through other classical genetic and embryological approaches. We generated an atlas of 489,686 single-cell transcriptomes from 62 closely spaced developmental stages spanning the first 5 days of zebrafish development, encompassing the first transcriptional events after zygotic genome activation to a freely-swimming, feeding animal. We used these data to search for gene expression programs that are reused across multiple tissues, determined the duration of transcriptional states during development, and identified dividing populations that are present for long developmental durations. Since global analysis of a dataset of this complexity rarely reveals its full cellular diversity, we also performed focused analyses within selected tissues. This enabled identification of uncharacterized cell subtypes and resolved the expression profiles of less characterized cell types within the intestine and non-skeletal muscle cell populations. For these less characterized cell types, we propose putative progenitors, candidate regulator genes, and cell type-specific gene expression cascades, using pseudotime trajectory approaches. To make these data accessible to the zebrafish community and other researchers, we developed the web portal, Daniocell.

## Results

### Temporally dense sampling of zebrafish development using single-cell RNAseq

Since developmental processes are incredibly dynamic, understanding them requires identifying the timing, ordering, and coordination of gene expression within and between cell types, and therefore requires assaying gene expression with high temporal resolution. Thus, to profile the molecular cell states that occur during wild-type zebrafish development, we generated single-cell transcriptomes from entire wild-type (TL/AB) zebrafish embryos and larvae across 50 closely spaced developmental stages, ranging from 14–120 hours post-fertilization (hpf). To capture underrepresented populations and allow robust cluster identification, we sampled heavily every 12 hours; to maintain temporal continuity necessary to investigate gene expression dynamics, we additionally sampled smaller cell numbers every 2 hours (Fig. 1a). In order to profile a large number of cells at an acceptable cost, we employed the cell hashing technique MULTI-seq (McGinnis et al. 2019), which barcodes cells prior to sample collection. Input cell concentrations were then increased, which profiled more cells, but at the cost of increased artefactual cell doublets (where multiple cells are incorrectly identified as ‘one cell’). The MULTI-seq hashing allowed computational identification and removal of doublets, which had multiple barcodes. We mapped reads to the GRCz11 genome, annotated using the Lawson Lab Zebrafish Transcriptome Annotation (v4.3.2) that harmonizes Ensembl and Refseq annotations, includes improved 3’ UTR models, and proposes additional gene models (Lawson et al. 2020). We also remapped published single-cell data from wildtype (TL/AB) zebrafish encompassing 3.3–12 hpf (Farrell et al. 2018) and combined the two datasets using Seurat to generate a continuous single-cell time-course of 489,686 cells from 62 closely-spaced developmental stages spanning 3.3 to 120 hpf (Fig. 1a– b, Supplementary Fig. 1a). Our dataset complements recent wild-type zebrafish single cell atlases that either use different techniques (single nuclei vs. single-cell), profile shorter developmental durations, or have lower frequency of collection timepoints (Farnsworth, Saunders, and Miller 2020; Farrell et al. 2018; Wagner et al. 2018; Saunders et al. 2022; Dorrity et al. 2022).

**Figure 1:**
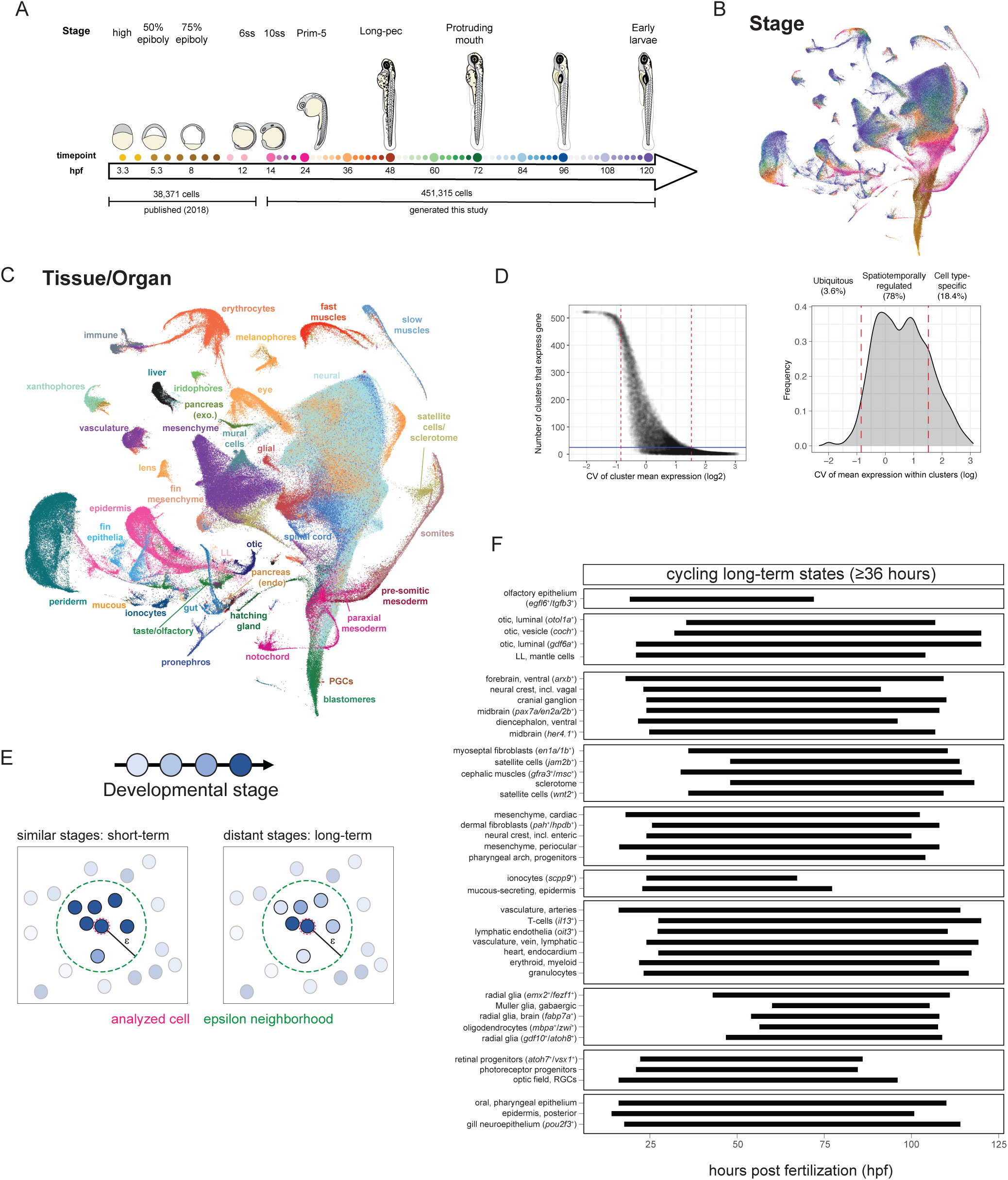
A high temporal resolution single-cell RNAseq timecourse encompassing embryogenesis and early larval development. (**A**) Single-cell transcriptomes were collected from whole zebrafish embryos at 50 different developmental stages (colored dots) between 14–120 hpf and then merged with our previous dataset encompassing 3.3–12 hpf (Farrell et al., 2018). Size of dots represents the number of cells recovered from each stage. (**B–C**) UMAP projection of single-cell transcriptomes, colored by (B) developmental stage (colored as in Fig. A) and (C) curated major tissues. **(D)** Distribution of the coefficient of variation (CV) of cluster means of expression for each gene (log-transformed). Lower CV indicates relatively similar expression across all clusters, while high CV indicates high variation in expression, either temporally or across cell types. Genes were divided into categories based on thresholds (red dashed lines). (**E**) Schematic showing our approach for identifying transcriptionally similar cells using an epsilon (ε) neighborhood approach and determining whether each cell was in a ‘short-term’ or ‘long-term’ state (based on the mean of absolute stage difference between the analyzed cell and its ε-neighbors). (**F**) Timeline bar plots showing the duration of ‘long-term’ cycling cell states identified using a 36-hour threshold. Each bar represents a cell population that was identified as ‘long-term,’ and the length of bar represents the minimum timespan that encompasses 80% of its ε-neighbors. hpf: hours post fertilization, PGCs: primordial germ cells, LL: lateral line.

We next identified transcriptionally distinct cell types and cell states during development (Fig.1c). In this work, we use the term ‘cell type’ to describe persistent cell identities (*e.g.,* muscle vs. blood), and ‘cell state’ to refer to more transient transcriptional profiles that cells enter and exit over time (*e.g.,* cycling vs. non-cycling or progenitor vs. differentiated). We iteratively clustered single-cell transcriptomes in a semi-supervised manner (see Materials and Methods), where we performed an initial clustering and assigned cells to 19 broad tissue types, based on cluster identity, transcriptional similarity, developmental stage, and established literature (Supplementary Fig. 1b). Then, within each tissue, we performed a second semi-supervised clustering, and clusters were annotated according to their expressed genes whose patterns were described by prior publications or the ZFIN gene expression database (Bradford et al. 2022). Altogether, this identified 521 clusters that represent ∼300 terminal and intermediate cell types or cell states. To visualize these data, cells were projected onto a Uniform Manifold Approximation and Projection (UMAP) and colored according to developmental stage and tissue of origin (Fig. 1b–c, Supplementary Fig. 1a–b). This dataset captures the specification and differentiation of cells comprising major organs, including the liver, gut, kidney, muscle, skin, circulatory system, and sensory systems.

To enable public access to our catalog of zebrafish developmental cell states from early embryogenesis to larval stages, we have created a static single-cell portal, Daniocell (https://daniocell.nichd.nih.gov/). This portal contains pre-computed information about gene expression during zebrafish development, rendered across all cells and each tissue separately. It is designed to answer the most common questions we expect researchers will ask from these data, namely: (1) When and where is any given gene expressed? (2) Which genes have the most similar and dissimilar gene expression patterns? (3) When is each cell type present and undergoing the cell cycle? and (4) Which genes does each cell type express most strongly and most specifically, and how does that compare to related cell types?

### Spatiotemporal variation of gene expression during development

We sought to determine for each gene in our transcriptome reference: (1) whether we captured its expression, and (2) whether its expression pattern was relatively constant or varied during development. While defining thresholds for these measures involves a degree of arbitrariness, they provide a rough measure of the comprehensiveness of these data and the portion of the genome that is deployed and regulated during development. We detected meaningful expression for 24,216 of the 36,250 genes in the transcriptome (67%), defined as ≥1 count in at least 22 cells (the size of our smallest cluster) and mean expression of ≥0.1 counts/cell in at least one cluster. To assay how many genes vary in expression across cell types and developmental stages, we used the coefficient of variation (CV) of cluster mean expression values as a rough measure. By relating CV to the number of clusters that express a gene (Fig. 1d) and by visually inspecting the expression pattern of randomly selected genes with different CVs (Supplementary Fig. 1c), we chose a set of thresholds to roughly categorize gene expression patterns. ∼3.6% of genes are ubiquitously expressed with low levels of fluctuation during development (logCV ≤ –0.85)—far fewer than identified by studies in adult organs (Fig. 1d) (Eisenberg and Levanon 2013; Hounkpe et al. 2021). ∼18% are ‘cell-type specific’ and expressed in ≤25 clusters, which generally corresponds to only a few cell types (logCV ≥ 1.52). The remaining 78% of genes exhibit spatiotemporal changes but are expressed in many cell types across several tissues. While scRNAseq does not capture all mRNA within each cell, these data do describe quantitative, temporally-resolved, and cell-type specific gene expression patterns for two-thirds of the genome during the first five days of zebrafish development. Moreover, it emphasizes that most genes vary spatiotemporally, suggesting that even general processes required for cellular survival vary between cell types or between developmental stages.

Additionally, about half of the genes annotated in the reference transcriptome we used lack proper gene names and have less experimental support. We measured whether they were detected with similar frequency and spatiotemporal variation, compared to ‘named’ genes. We categorized genes into four groups based on their level of evidence: ‘named’ genes (54%), unnamed cDNA clones (e.g. genes beginning with *si:ch211-*, *si:dkey-*, *zgc:*, etc., 13%), computationally predicted genes (e.g. genes beginning with LOC-, CABZ-, BX-, AL-, etc., 27%), and putative genes identified through *de novo* transcriptome assembly by the Lawson lab (genes beginning with XLOC-, 5.4%). Unnamed cDNA clones were detected with similar frequency to ‘named’ genes (67%) and exhibited spatiotemporally varying expression with slightly higher frequency (Supplementary Fig. 1d). Computationally predicted and ‘XLOC–’ genes were detected less frequently (28% and 37% respectively), but when detected were more likely to be spatiotemporally restricted, lending additional confidence that they may represent uncharacterized genes. Altogether, these results suggest that many of these unstudied genes are indeed expressed and regulated during development. Given their frequent sequence similarity to other ‘named’ genes, it is possible that many provide redundant functions and may have masked the roles of some important developmental regulators from forward genetic screens.

### Mapping the duration of transcriptional states using a genome-wide approach

Developing cells transition through several transcriptional states during specification and differentiation. Our dataset provides an opportunity to use cells’ full transcriptomes to identify the duration of each transcriptional state (i.e., how long cells in that transcriptional state can be found during development). Transcriptional states that are present for short durations occur only at a specific timepoint during development. Other transcriptional states that are present for longer may indicate that individual cells are persistently in this state for a long period of time (for instance, if a terminally differentiated cell adopts a transcriptome that is stable over time), or may represent a state that individual cells occupy for a short time, but during many developmental stages (for instance, in an asynchronous developmental process like somitogenesis, or if a similar transcriptional state is used by multiple cell types at different times). Using our dataset, we sought to (1) determine the duration of transcriptional states during development, (2) assess whether cell populations were undergoing continued cell division, and (3) use these combined analyses to identify potential long-term progenitor states.

To identify the duration of developmental states, we used an ε-nearest neighbor approach: we (1) defined a distance in gene expression space to represent transcriptionally similar cells (ε), (2) found each cell’s neighbors within that ε-neighborhood, and (3) computed the difference in developmental stage between each cell and its neighbors (Fig. 1e). Cells were categorized based on the mean of absolute stage difference with their neighbors (e.g., <24h, 24–36h, 36–48h, >48h) (Supplementary. Fig. 2a–d). Cells in transcriptional states that are present “long-term” would be expected to have neighbors that were more different in developmental stage than cells in transcriptional states that occur only during a limited period of development, whose neighbors should all have a similar stage.

Most transcriptional states during development were present for a limited time. During the first 5 days, 18% of cells were in transcriptional states that last for ≤12 hours, 58% that last for ≤24 hours, 81% that last for ≤36 hours, 93% that last for ≤48 hours, and only 7% of cells were in a transcriptional state found for >48 hours. For downstream analyses, we classified cells into “short-term” and “long-term” states based on a threshold of 36 hours—transcriptional states present ≥36 hours were considered “long-term” (Supplementary Fig. 2e). This threshold was chosen to balance focusing on states whose duration was rare while avoiding approaching the upper limit possible for this analysis on a time course of 120 hours. Additionally, each cell was classified as “cycling” or “non-cycling” based on its expression of transcripts associated with different cell cycle phases (Supplementary Fig. 2, Supplementary Fig. 3).

During terminal differentiation, many cell types exit the cell cycle and adopt a stable transcriptome. Correspondingly, we found that most cells in “long-term” transcriptional states had exited the cell cycle (68% of ‘long-term’ cells were non-cycling), and these cells were generally from later developmental stages (74% were 72 hpf or older); we interpret these as terminally differentiated. Interestingly, certain tissues predominantly exhibited “short-term” transcriptional states even when cells exited the cell cycle during later stages, including the periderm (76% of non-cycling cells in “short-term” states), fin (85%), eye (88%), and endoderm (94%) (Supplementary Fig. 3). This suggests that in these tissues, even after cells stop dividing, differentiate, and become functionally required, their transcriptional profile continues to change. For the endoderm, this result aligns with previous observations that zebrafish digestive organs (liver, pancreas, and intestine) mature in larvae after 96 hpf, although some of these organs become physiologically functional as early as 76 hpf (Li et al. 2020; Wallace et al. 2005; Flores et al. 2020).

The remaining 32% of cells in “long-term” transcriptional states were still undergoing cell division. Some of these cells are in non-differentiated developmental states that exhibit high rates of cell division and that cells enter over a prolonged period due to developmental asynchrony; for instance, we recover populations of transit amplifying cells like primitive erythroblasts, myoblasts, and periderm progenitors. Alternatively, some of these cells represent known long-term stem cells—such as radial glia, muscle satellite cells, hematopoietic stem cells, mesenchymal progenitors, and some neural progenitors (e.g. *her4.1^+^*) (Fig. 1f, Supplementary Fig. 2e). This suggests that these known progenitor populations remain transcriptionally consistent at the whole transcriptome level over a long duration. Recent work in zebrafish neuronal development similarly highlighted that some retinal progenitors remain transcriptionally consistent—there are retinal progenitors at 15 dpf that are nearly transcriptionally identical to embryonic progenitors that appear prior to 24 hpf (Raj et al. 2020). Here, we find that this is a common theme among eye progenitor populations, and several progenitor populations that have been studied with individual marker genes (Nelson, Park, and Stenkamp 2009; Chuang and Raymond 2001; Kennedy et al. 2004; Bando et al. 2020; Shen and Raymond 2004) are found “long-term” even when considered at a whole-transcriptome level, including (i) *rx1*^+^/*rx2*^+^/*rx3*^−^ and *rx3*^+^/*otx2*^+^ optic progenitors, (ii) photoreceptor progenitors, and (iii) progenitors of retinal ganglion cells, cone bipolar cells and oligodendrocytes (*atoh7*^+^/*vsx1^+^*/*olig2*^+^) (Fig. 1f). These results contrast with some other classic stem cell populations that mature transcriptionally during development — for instance, *pax3*+ spinal cord progenitors are present from 14–58 hpf, but their full transcriptional profiles differ markedly over that time, and they exhibit a transcriptional persistence of <24hpf. This has been similarly shown previously for *insm1a/her4.1+* hypothalamic progenitors (Raj et al. 2020).

While this analysis recovered many known stem cell populations, it also identified some cycling populations as “long-term” that are not well characterized: (i) an *oit3*+/*lyve1a*^+^ lymphatic population, (ii) cephalic muscles (*gfra3*^+^/*ret*^+^/*msc*^+^), (iii) dermal fibroblasts (*pah*^+^/*hpdb*^+^) in the pharyngeal arches, and (iv) myoseptal fibroblasts (*en1a/en1b*^+^/*vegfc*^+^) (Fig. 1f). These may represent unappreciated stem cell states or transit amplifying states and merit future investigation.

This analysis has some caveats. For instance, a “long-term” transcriptional state that appeared in the last 36 hours of the timecourse (84–120 hpf) could be classified as “short-term” because we lack measurements after 120 hpf. Additionally, in a few cases, progenitor populations are identified as “long-term” because of strong transcriptional resemblance between the dividing progenitors and their non-dividing, differentiated counterparts. Examples include early erythroblasts, which begin to express differentiation genes, such as hemoglobins (e.g. *hbbe1.1*, *hbbe1.2*) while they are still dividing and also expressing genes associated with progenitor states (e.g. *drl*, *blf*, *tal1*, and *urod*). Similarly, some periderm progenitors begin to express transmembrane and adhesion-related differentiation genes (e.g. *lye*, *anxa1a*, *krt17*, *eppk1*, *pkp1b*, *tm9sf5*) while still cycling. This results in smaller-than-ε transcriptional differences between these dividing progenitor and non-dividing differentiating cells and the subsequent overestimation of developmental duration of the progenitor state. We have excluded such examples from our analyses above. However, most progenitor populations only identify other progenitors as ‘transcriptionally similar’, thereby enabling discrimination between the two progenitor strategies described above.

Altogether, our analysis identified that most transcriptional states during the first 5 days of zebrafish development are present for ≤36 hours. We find that many cell types adopt a stable transcriptome as they differentiate, but we highlight that this varies between tissues. Additionally, we captured multiple populations of “long-term” cycling cells. This included many known stem cell populations and thereby identifies which stem cells are more likely to have a stable transcriptome during early development. It also highlighted a few “long-term” transcriptional states as candidate unstudied stem cell populations.

### Identification of shared developmental gene expression programs and tissue-specific adaptations

Many distinct cell types share common cellular states, features, or elaborations. For example, despite providing dramatically different functions, olfactory sensory neurons and kidney tubular cells both have cilia. While ciliogenesis is known to result from a shared genetic program, for many such shared features, it remains unclear whether the underlying genetic program is also shared. Whole-animal, time-course, single-cell RNAseq data can help address this question, as it enables gene expression correlation analysis across all cell types and many developmental stages. Here we catalogued gene expression programs (GEPs) that are shared between two or more tissues during zebrafish development (Fig. 2a). To find these GEPs, we: (1) smoothed the data from all cells in our dataset using a 5 nearest-neighbor network to reduce effects of technical noise (Wagner, Yan, and Yanai 2018), (2) used fuzzy c-means (FCM) clustering to group genes with similar expression over time and across tissues (Hall et al. 1992; Cannon, Dave, and Bezdek 1986; Bezdek 1980; Kumar and M 2007), (3) filtered out poor quality GEPs (<5 member genes or primarily technical member genes), and (4) filtered out GEPs that were expressed in a single tissue. As a complementary approach, we also calculated GEPs on tissue-specific subsets and used correlation with cosine distance to identify modules that were found in multiple tissues; while this approach identified many more cell-type specific GEPs, it did not yield any additional shared GEPs that were not already captured from the global analysis (data not shown). This approach identified 90 GEPs shared between multiple tissues, comprised of 8–735 member genes (average 95 member genes). We were able to clearly annotate 82 of these shared GEPs based on known functions of individual member genes (Fig. 2a, Supplementary Table 2); it is currently unclear whether the remaining 8 represent technical artifacts or novel GEPs without description in the literature.

**Figure 2:**
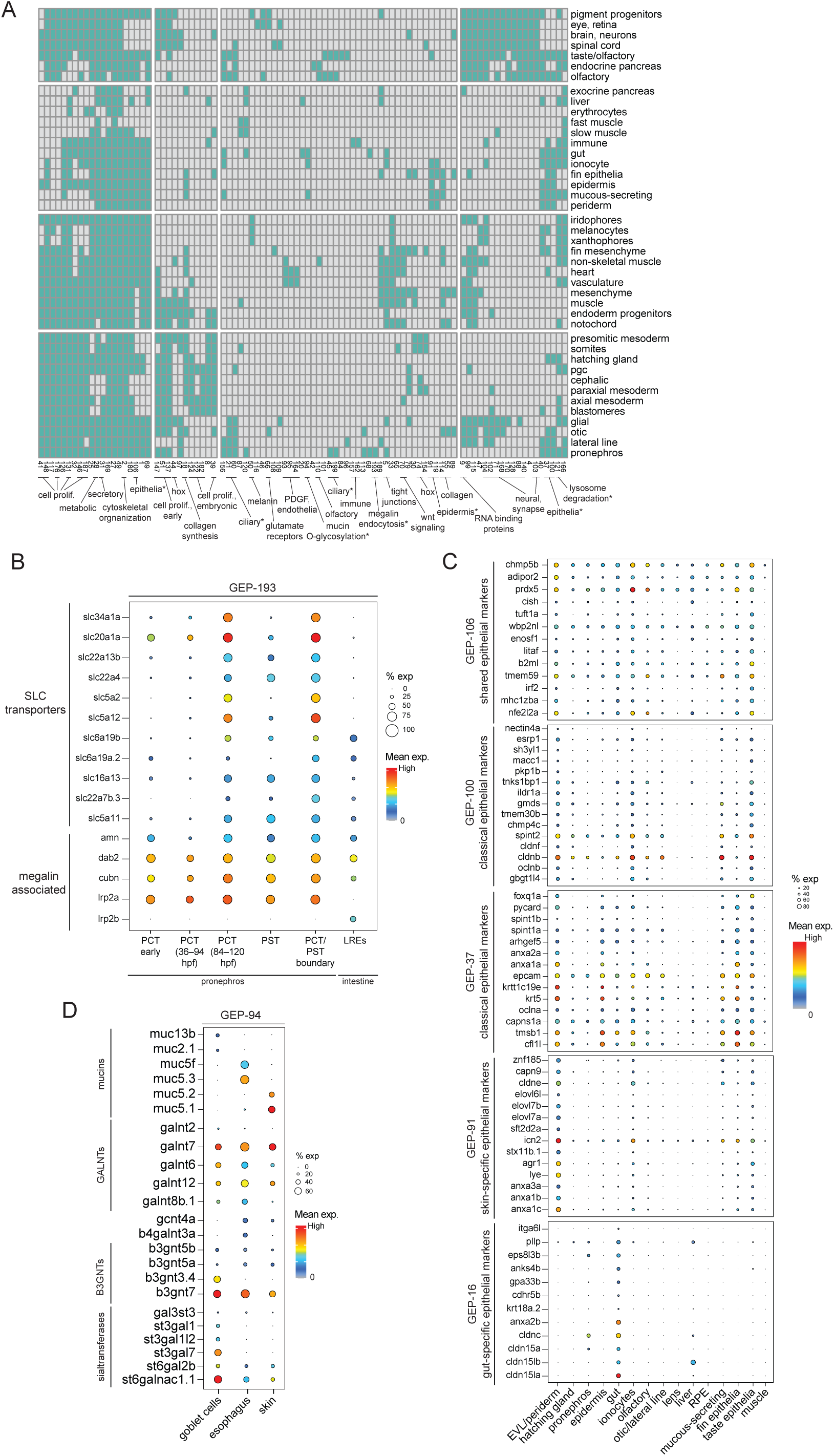
Identification of gene expression programs shared by multiple distinct cell types during development. (**A**) Binary heatmap showing expression domains of gene expression programs (“GEPs”, x-axis) reused in multiple tissues (y-axis) during development. Annotations for select GEPs are shown. Asterisk indicates the GEPs further investigated in panels B, C, D, and Supplementary Fig. 5. Full GEP annotation can be found in Supplementary Table 2. (**B**) Dot plot showing expression of Megalin-associated genes and SLC transporters from module GEP-193 that are shared between intestinal lysosome-rich enterocytes (LREs) and pronephros proximal convoluted tubule (PCT) and pronephros proximal straight tubule (PST) cells. (**C**) Dot plot of top-loaded genes from five epithelial GEPs: one shared universally across all epithelial cell types (GEP-106), two comprising classical epithelial genes (GEP-100, GEP-37), and two tissue-specific (GEP-91, GEP-16). X-axis: cell types with epithelial characteristics across several tissues, and muscle as a non-epithelial control. (**D**) Dot plot of shared and tissue-specific members of module GEP-94, associated with mucin O-glycosylation. For panels B–D, color: mean expression per cell type; size: percent of cluster cells that express each gene on the Y-axis. **B3GNTs**: β1,3-N-acetylglucosaminyltransferases; **GALNTs**: N-acetylgalactosaminyltransferases; **RPE**: retinal pigmented epithelium; **EVL**: enveloping layer.

Out of the 82 annotated GEPs, 20 represented ubiquitous GEPs shared across most or all tissues, involved in essential cell functions such as metabolism, cell cycle, or cytoskeletal organization (Fig. 2a, Supplementary Table 2). The remaining 62 GEPs were restricted to particular cell types across 2 or more tissues and generally represented functionally related genes involved in conferring specific cellular features. For instance, as a confirmation of our approach, we identified four GEPs associated with one of the most well-studied recurring cellular features—the motile cilium (Supplementary Fig. 4a). Three modules (GEP-21, GEP-45, GEP-60) contained genes involved in motile cilia assembly, while the fourth (GEP-199) comprised genes expressed specifically in multi-ciliated cells (Marra et al. 2019; Zhou et al. 2017).

In some cases, modules demonstrated cellular features that have a core, shared network that is then modified within different cell types. For instance, we observed co-expression of Megalin-associated receptor-mediated endocytic machinery genes that facilitate protein absorption (GEP-193: including *cubn*, *amn*, *dab2*), which was known to be shared between intestinal lysosome-rich enterocytes (LREs) and pronephros proximal tubules (Zhang et al. 2013) (Fig. 2b). *megalin* itself was previously only documented in the pronephros (Park et al. 2019), but GEP-193 identified both paralogs: *lrp2a* as pronephros-specific and *lrp2b* as intestine-specific (Fig. 2b). This module also included several membrane-spanning transporters that transport sodium/glucose (*slc5a11*), organic ions (*slc22a7b.3*), monocarboxylate (*slc16a13*), and amino acids (*slc6a19a.2*, *slc6a19b*), highlighting that Megalin-associated endocytosis is a component in a broader process for nutrient reabsorption (Fig. 2b). Moreover, these tissues additionally expressed module GEP-121 which contained lysosome-associated catabolic enzymes, including proteinases, hexosaminidases, and mannosidases (Supplementary Fig. 4b). These genes have previously been associated only with intestinal LREs (Wen et al. 2021), but this shows that increased protein reabsorption via Megalin-associated endocytosis is accompanied by a corresponding increase in lysosomal degradation capacity in other tissues as well. Despite these core similarities, these modules also captured tissue-specific adaptations to these processes. For instance, GEP-121 was also shared with other lysosome-rich cell types that are not involved in nutrient reabsorption, such as melanophores, macrophages, microglia, and lymphatic endothelia, indicating that despite co-expression in some cell types, lysosomal degradation capacity and protein reabsorption activity are separately regulated (Supplementary Fig. 4b). Additionally, GEP-193 included several pronephros-specific transporters for sodium/glucose (*slc5a2*, *slc5a12*), organic ions (*slc22a4*, *slc22a13b*), and phosphate transporters (*slc20a1a*, *slc34a1a*), highlighting that the shared reabsorption program is further specialized in individual tissues (Fig. 2b).

Similar themes were observed in gene expression programs that conferred cellular features that were shared even more broadly. For instance, we observed five shared GEPs across epithelial cell types (Fig. 2c), including two (GEP-100 and GEP-37) comprised of traditional epithelial marker genes (e.g. *epcam*, annexins, occludins, claudins, and keratins). Interestingly, those ‘traditional’ epithelial markers were excluded from some epithelial cell types, such as the lens, liver, and retinal pigmented epithelium (RPE). However, we identified GEP-106 that was shared across all epithelial cell types (including cell types that express ‘traditional’ markers and ones that do not) which included *adiponectin receptor 2* (*adipor2*), the acidic phosphorylated glycoprotein (*tuftelin*/*tuft1a*), MHC-class I antigen (*mhc1zba*), *beta-microglobulin* (*b2ml*), *enolase superfamily member 1* (*enosf1*), and a bZIP transcription factor (*nfe2l2a*). This suggests a less-explored core gene expression program that may influence epithelial biology in broader epithelial types, including even non-canonical epithelia. This network may then be modified by expression of additional programs, such as the ‘traditional’ epithelial markers or one of two tissue-specific epithelial signatures also identified here – GEP-91 in the periderm, ionocytes, and mucous-secreting cells, and GEP-16 in gut epithelia (Fig. 2c). Similarly, we found a core, shared expression program across mucous-secreting cells within the intestine, esophagus, and skin (GEP-94). In addition to mucins (a large family of heavily O-glycosylated proteins), this module identified a group of O-glycosylation-catalyzing enzymes which were shared between all mucous-producing cell types and expressed with nearly identical dynamics in those cell types (Fig. 2d, Supplementary Fig. 5), including GALNTs (which add N-acetyl galactosamine to the mucin backbone) and B3GNTs and B4GNTs (which elongate the glycan chain by adding galactose). However, intestinal goblet-cells expressed more sialtransferases (that adds sialic acid as terminal sugars) than esophagus or skin mucous-secreting cells, indicating that goblet-cell mucins may have longer sialic acid chains. This suggests that differential mucin modifications in zebrafish may mirror those in other animals, such as rat, where longer sialic acid chains have been hypothesized to protect gut mucins from bacterial proteolytic enzymes (Grondin et al. 2020).

Altogether, these results highlight that while some cellular features are produced by re-using gene expression programs during development across multiple tissues, the catalog is actually relatively limited, and that in most cases, those shared programs are accompanied by cell-type-specific elaborations that customize them for each distinct cell type.

### Focused analysis of zebrafish non-skeletal muscle identifies new pericyte subpopulations

An immense anatomical cell type diversity is present during zebrafish development, and iterative clustering within this whole-animal dataset revealed a similar transcriptional diversity. However, there is not a perfect correspondence between these two categorizations. So, we performed focused analyses within non-skeletal muscle cells and endodermal derivatives to further explore their transcriptional diversity and to better align it with functional and anatomical categorizations.

Non-skeletal muscle primarily consists of smooth muscle cells (SMCs), which are a type of involuntary muscle that form interconnected sheets and provide structural and functional support to luminal organs, including the digestive system (visceral smooth muscle) and the circulatory system (vascular smooth muscle). In addition to vascular smooth muscle cells (vaSMCs), which surround major blood vessels, the circulatory system is also supported by pericytes, which surround capillaries. Pericytes and vaSMCs are spatially and morphologically distinct, since pericytes associate with different vessels, exhibit cytoplasmic extensions or projections, and are scattered along capillaries, rather than forming sheets (Dore-Duffy and Cleary 2011; Hartmann et al. 2015). However, these two cell types exhibit considerable transcriptional and functional overlap, since both vaSMCs and pericytes can control blood pressure, sculpt neighboring epithelia, and regulate neighboring tissue by producing secreted signals (Stratman et al. 2017; Smyth et al. 2018; Yamazaki and Mukouyama 2018; Baek et al. 2022). However, the transcriptional heterogeneity, spatial distribution, and developmental regulators of pericytes remain areas of active investigation in multiple organisms (Ando, Ishii, and Fukuhara 2021; Donadon and Santoro 2021). To further explore cellular heterogeneity and gene expression profiles within these enigmatic tissues in zebrafish, we iteratively re-clustered 3,866 non-skeletal muscle cells (which include smooth muscle and pericytes) (Fig. 3a, Supplementary Fig. 6a). Our clusters include 2 cardiac muscle populations (clusters C14 and C17), hepatic stellate cells (C18), and 2 *pdgfra*+ populations that we were unable to annotate (*lyve1a*+ or *cxcl11*+, C5 and C19). We identified smooth muscle based on expression of the traditional smooth muscle markers *acta2* and *tagln*, which included five vascular SMC populations (C2, C11, C12, C15, and C21). Additionally, we identified three visceral SMC populations (C8, C10, C13) based on combined expression of the traditional visceral SMC markers *desmin-b* (*desmb*) and *smoothelin-a and b* (*smtna* and *smtnb*) (Kayman Kurekci et al. 2021; Georgijevic et al. 2007) (Fig. 3b). We also identified three potentially visceral SMC populations (C16, C22, and C23) that expressed only one or two of those three traditional visceral SMC markers (Supplementary Fig. 6b). Lastly, we identified three transcriptionally distinct pericyte populations (C4, C9, and C20) and a population of myofibroblasts (C3) (Fig. 3c, Supplementary Fig. 6c). Vascular SMCs are relatively well characterized, so in this section, we focus on the distinct pericyte populations and return to visceral SMCs in the next section.

**Figure 3:**
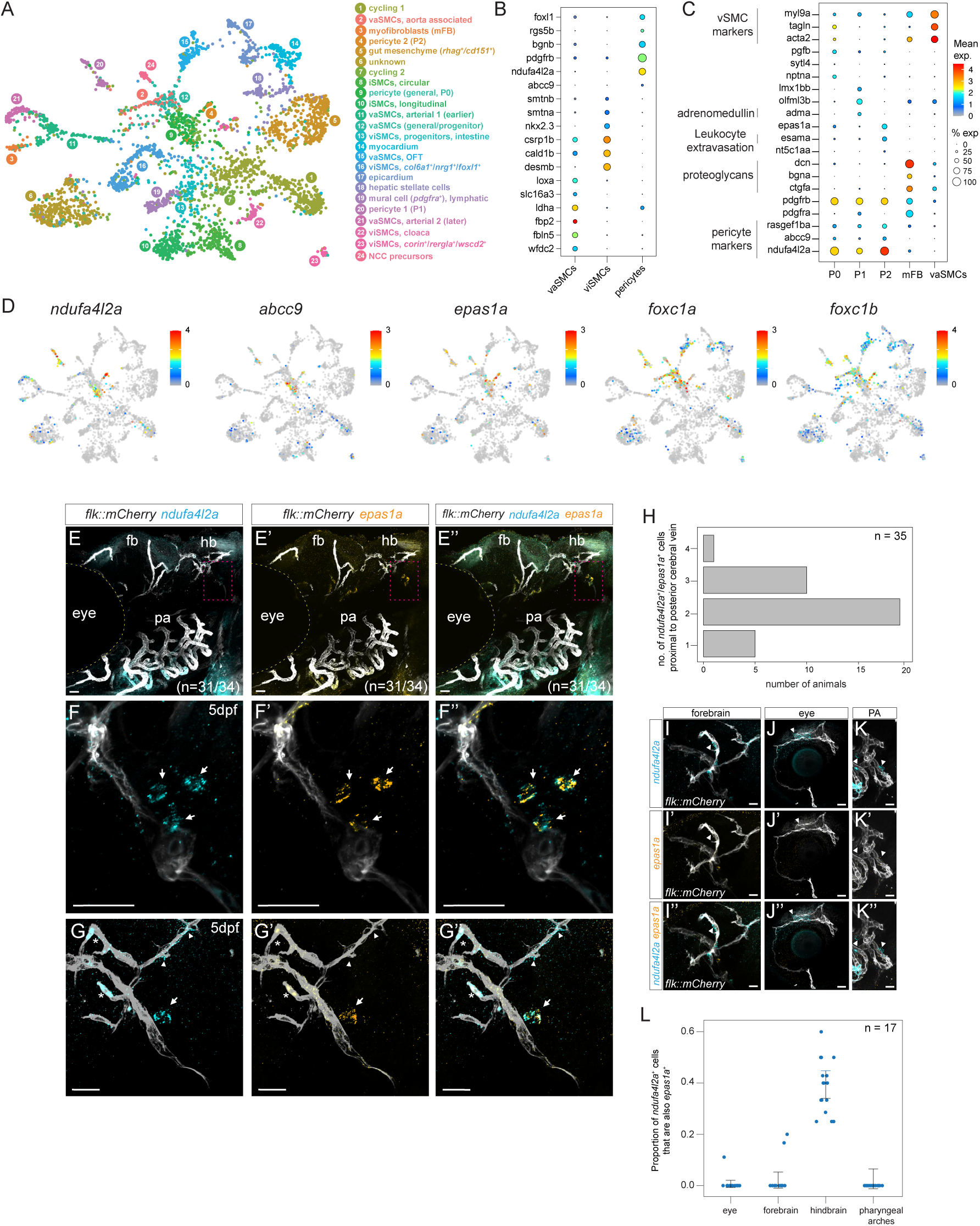
Sub-clustering of non-skeletal muscle reveals distinct pericyte subtypes. (**A**) UMAP projection of 3,866 non-skeletal muscle cells, numbered and color-coded by cluster. (**B**) Dot plot of selected differentially expressed pericyte-specific markers (x-axis) compared to vascular (vaSMCs) and visceral SMCs (viSMCs). **(C)** Dot plot of selected differentially expressed genes (y-axis) between the three pericyte clusters (x-axis, P0–P2) and myofibroblasts compared to vascular SMCs (x-axis, vaSMCs). See Supplementary Fig. 6C for additional markers. (**D**) Expression of pericyte marker genes visualized on the UMAP projection. Color bar shows mean expression level of each gene. (**E–G’’**) RNA *in situ* hybridization for markers specific to the pericyte-2 population on a *flk::mCherry-CAAX* background in 5 dpf larvae. Panels E–E’’ indicate lateral view of the whole zebrafish head for the universal pericyte marker *ndufa4l2a* and pericyte-2 marker *epas1a*. Panels F–G’’ indicate a higher magnification of the brain posterior cerebral vein with *epas1a*^+^ pericytes. Arrows indicate *ndufa4l2a*^+^/*epas1a*^+^ pericytes in contact with as well separated from the posterior cerebral vein. In panels G– G”, arrowheads indicate *ndufa4l2a*^+^/*epas1a*^−^ pericytes along other hindbrain vessels while asterisks indicate autofluorescent red blood cells inside blood vessels. (**H**) Bar graph quantifying the number of *ndufa4l2a*^+^/*epas1a*^+^ pericytes that were visible in a similar-sized field of view near the posterior cerebral vein, per animal. (**I–K’’**) RNA *in situ* hybridization for *ndufa4l2a* and *epas1a* across other blood vessels in the zebrafish head including forebrain (I–I”), eye (J–J”), and pharyngeal arches (K–K”). Arrowheads mark cells that are *ndufa4l2a*^+^; no *ndufa4l2a*^+^/*epas1a*^+^ cells were observed in these regions. (**L**) Jitter plot showing the proportion of *ndufa4l2a*^+^ pericytes that were also *epas1a*^+^ in different regions of the zebrafish head. Error bars indicate standard error of mean (S.E.M). **PA**: pharyngeal arches. Scale bar: 25 µm.

Pericyte clusters were identified based on the expression of previously described marker genes in mouse and zebrafish such as *abcc9*, *pdgfrb* and the recently described pericyte-specific (within zebrafish perivascular cells) marker gene, *ndufa4l2a* (Shih et al. 2021; Vanlandewijck et al. 2018)(Fig. 3c–d, Supplementary Fig. 6c, Supplementary Fig. 7a). Transcriptionally distinct pericyte subpopulations have been previously demonstrated in mouse (Vanlandewijck et al. 2018) but have not been described in zebrafish. Here, we observed three distinct transcriptional states or subpopulations of pericytes – pericyte-0 which does not have any unique markers, and two others with at least 10 specific markers (Supplementary Fig. 6c). As examples of these specific markers, pericyte-1 expresses *adrenomedullin* (*adma*), and pericyte-2 expresses leukocyte extravasation genes such as *esama* and *epas1a* (Fig. 3c–d). A few cells within the pericyte-1 cluster also expressed the characteristic genes of pericyte-2, which indicates that the two subpopulations may be closely related or represent non-exclusive transcriptional states (Fig. 3c). Expression of some traditional pericyte and SMC markers varied among these populations. For instance, *pdgfra* (often considered a fibroblast marker) was expressed in pericyte-1, and the traditional SMC marker *tagln* was expressed in pericyte-0, but excluded from pericyte-1 and pericyte-2 (Fig. 3c). Previous studies have described that pericytes originate from multiple embryonic origins, including the sclerotome, neural crest, and lateral plate mesoderm (Le Lievre and Le Douarin 1975; Jiang et al. 2000; Pouget et al. 2006; Etchevers et al. 2001; Korn, Christ, and Kurz 2002). Cranial pericytes derive from the cranial neural crest in zebrafish and mammals (Whitesell et al. 2014; Ando et al. 2016), and at the developmental stages profiled, most pericytes in zebrafish are located in the head. Consistent with this, all three pericyte clusters expressed the classical cranial neural crest marker *foxc1a/b* (Whitesell et al. 2019; French et al. 2014), which suggests that these populations derive from the neural crest (Fig. 3d).

Additionally, we observed a fourth population (C3) that expressed *acta2*, *pdgfrb*, and *ndufa4l2a* at lower levels, but also expressed proteoglycans (*bgna*, *ctgfa* and *dcn)* and ECM components (including *col6a1/2*, *col4a5/6, and col11a1a*) that are traditionally associated with fibroblasts (Muhl et al. 2020) (Fig. 3a, c, Supplementary Fig. 6c, Supplementary Fig. 7). Based on its gene expression and its localization far from vessels (Supplementary Fig. 7), we believe this population represents myofibroblasts. However, given its weak expression of the pericyte marker *ndufa4l2a* and documented transdifferentiation between pericytes and fibroblasts (Sundberg et al. 1996), it remains possible that these cells have a more complicated identity.

A potential artifactual explanation for observing multiple pericyte signatures would be that pericytes incompletely dissociated from neighboring cells and formed doublets with other endothelial or mesenchymal cells. To exclude this possibility, we simulated incomplete dissociation by creating artificial doublets between pericyte-0 cells and other cell types that expressed the characteristic markers of pericyte-1 and pericyte-2 subtypes (see Methods). None of these artificial doublets recreated the signatures observed in the pericyte-1 and pericyte-2 populations (Supplementary Fig. 8a– b), suggesting that they do not represent artifacts generated by scRNAseq, but are real cellular states.

To further characterize the pericyte-2 population, we performed *in situ* hybridization for the pericyte-2 specific marker, *epas1a*, in combination with the general pericyte markers *ndufa4l2a* and *abcc9*. We observed *epas1a*+/*ndufa4l2a*+ and *epas1a*+/*abcc9*+ cells surrounding the posterior cerebral vein in the hindbrain (Fig 3e–g, arrows, Supplementary Fig. 8c). *ndufa4l2a*+ cells surrounding other hindbrain vessels did not express *epas1a*, suggesting that this indeed represents transcriptional heterogeneity (Fig. 3g, arrowheads). Interestingly, *ndufa4l2a*+/*epas1a*+ cells were frequently observed in pairs (Fig. 3f, h), and though *ndufa4l2a*+ cells were generally spatially proximal to vessels, for pairs of *ndufa4l2a*+/*epas1a*+ cells, often one cell of the pair was separated from the vessel. Inspection of other regions of the zebrafish head, including the forebrain, eye, and pharyngeal arches revealed many *ndufa4l2a*+ cells, but almost none that co-stained with *epas1a*, suggesting that the pericyte-2 population is spatially restricted to the hindbrain (Fig. 3i–l). This mirrors other organisms, where transcriptionally distinct pericyte subpopulations have been associated with particular tissues (Vanlandewijck et al. 2018). Altogether, these results reveal that there are multiple transcriptionally distinct pericytes in zebrafish that exhibit spatial segregation, much like mammals.

### Molecular characterization of zebrafish intestinal smooth muscle cell types

The intestine is surrounded by smooth muscle that regulates its morphogenesis (Huycke et al. 2019), stem cell maintenance (Martin-Alonso et al. 2021), enteric nervous system patterning (Graham et al. 2017), and gastrointestinal motility (Wedel et al. 2006; Bitar 2003). The intestinal smooth muscle (a subtype of visceral SMCs) is arranged in two layers, one oriented longitudinally and the other circularly (Georgijevic et al. 2007). In humans, mice, and zebrafish, intestinal SMC markers that label both circular and longitudinal smooth muscle have been reported (Sanders et al. 2012; Huycke et al. 2019; Martin-Alonso et al. 2021; Ma et al. 2018); however, to our knowledge markers distinguishing between circular and longitudinal SMCs have remained undescribed. Our focused analysis within non-skeletal muscle revealed two transcriptionally distinct *desmb*^+^/*smtnb*^+^ intestinal SMC populations in zebrafish with distinct markers. C8 expressed *kcnk18*, *fsta*, *foxf2a*, *gucy1a1*, *npnt*, and C10 expressed *il13ra2*, *tesca*, *rgs2*, *fhl3b* (Fig. 3a, Fig. 4a–b)*. In situ* hybridization for the C8/C10 markers *il13ra2*, *fsta*, and *kcnk18* identified C8 as the inner, circular layer of intestinal SMCs and C10 as the outer, longitudinal layer of intestinal SMCs (Fig. 4c–m, Supplementary Fig. 9a–h), demonstrating the distinct molecular profiles of the two layers for the first time.

**Figure 4:**
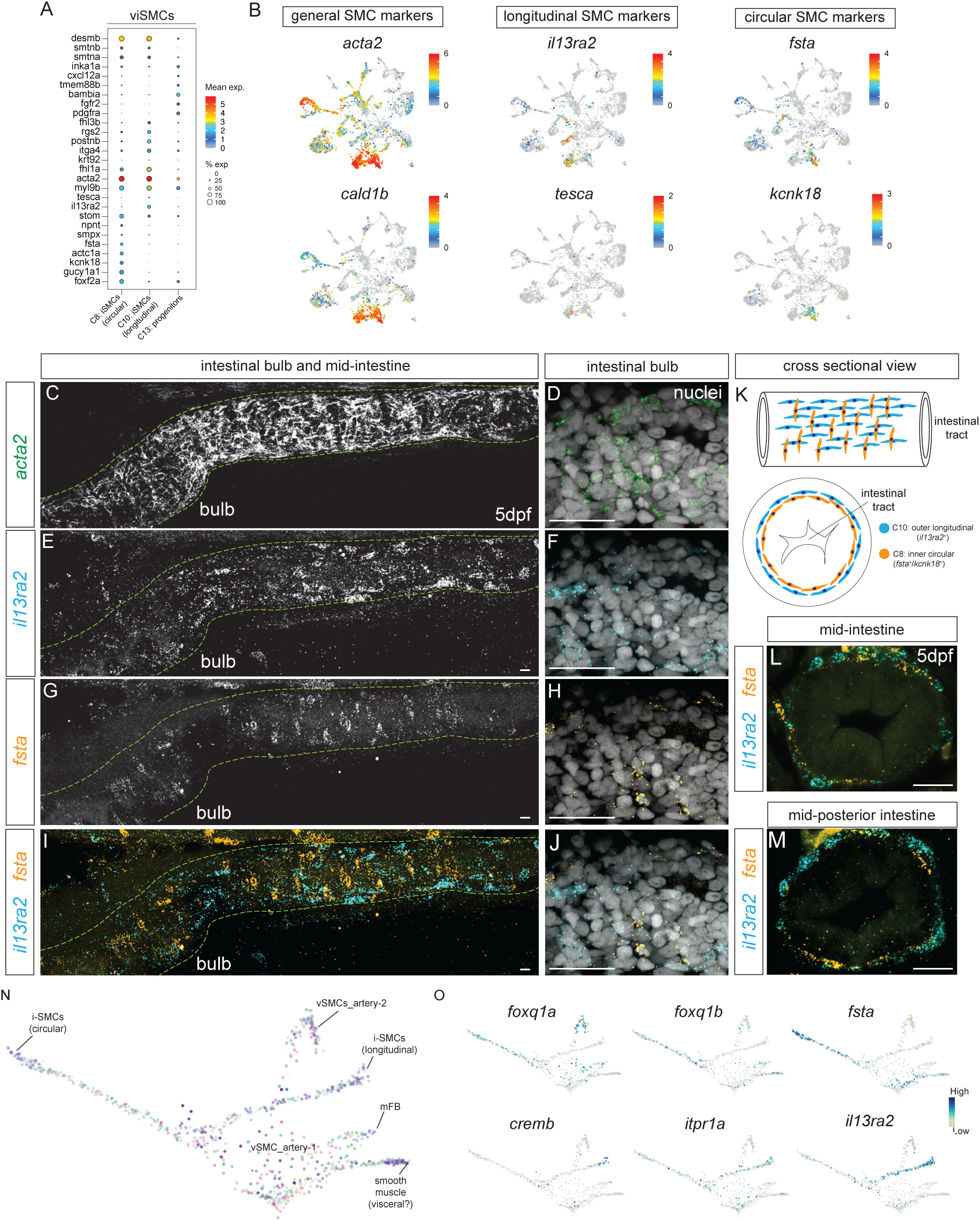
Distinct gene expression within longitudinal and circular intestinal smooth muscle cells. (**A**) Dot plot of top differentially expressed genes (y-axis) between circular and longitudinal SMC clusters (x-axis). (**B–D**) Feature plots of common (*acta2*, *cald1b*) and differentially expressed (*il13ra2*, *tesca*, *fsta*, *kcnk18*) markers of intestinal SMCs. (**C–J**) RNA *in situ* hybridization of markers expressed in longitudinal (*il13ra2*) and/or circular smooth muscle (*fsta*) cells lining the zebrafish intestinal tract. (C, E, G, I) anterior intestine view, including the intestinal bulb. (D, F, H, J) Higher magnification images of the intestinal bulb with nuclear (Hoecsht 33322) counterstain showing spatially segregated association of *il13ra2* and *fsta* with differently oriented nuclei. **(K)** Diagram showing the orientation of circular and longitudinal SMCs along the intestinal tract from a lateral view (top) and cross-section (bottom). (**L–M**) Expression of longitudinal (*il13ra2*) and circular (*fsta*) SMC-specific markers shown from a cross-sectional view. **(N)** Force-directed layout of an URD-inferred hierarchical tree calculated on *foxc1a/b*^−^ and *prrx1a/b*^−^ (putatively non-neural crest derived) pericytes and smooth muscle cells. The trajectory tree is colored by stage, as in Fig. 1A. (**O**) Expression of intestinal smooth muscle layer-specific TFs and markers visualized on the URD trajectory. Scale bar: 25 µm.

To identify the transcriptional events accompanying the acquisition of intestinal circular and longitudinal SMC fates, we reconstructed developmental trajectories using URD (Farrell et al. 2018). We focused on cell populations from the non-skeletal muscle atlas that were putatively non-neural crest derived (*foxc1a/b*– and *prrx1a/b*–) and expressed the general smooth muscle marker *acta2*, which included three visceral SMC populations, two vascular SMC populations, and the putative myofibroblasts. We reconstructed trajectories from 3 dpf progenitors to distinct 5 dpf clusters, which predicted a close relationship between the two types of intestinal SMCs (Fig. 4n). To uncover the distinct gene expression cascades that characterize these two layers of smooth muscle and to predict potential transcriptional regulators within them, we examined gene expression dynamics along the two intestinal SMC branches, identifying markers specific for each SMC including early onset transcription factors (TFs) that may be important for the specification of these two populations (Fig. 4o, Supplementary Fig. 9i–k, Supplementary Fig. 10). Several TFs were shared between the two intestinal SMC populations, including *foxf1*, *foxp4*, *meis2a*, and *pbx3b* (Supplementary Fig. 9i), though these factors were each expressed in numerous other cell types in the animal. However, we found that expression of the TFs *foxq1a*, *foxq1b, foxf2a* and *tcf21* was restricted to the circular SMC trajectory, while the longitudinal SMC cells instead expressed *cremb*, *itpr1a*, and *tead3a* (Fig. 4o, Supplementary Fig. 9j–k). Interestingly, despite their widespread expression, previous studies in mouse (Hoggatt et al. 2013) and *Xenopus* (Tseng, Shah, and Jamrich 2004) have demonstrated a requirement for *foxf1* and *foxf2* for general intestinal SMC differentiation. Altogether, these results revealed transcriptional differences between these two intestinal smooth muscle layers that may underlie some of their described morphological and biophysical differences (Huycke et al. 2019).

### Focused analysis of endodermal development identifies disease-relevant cell types and their candidate regulators

Similar to non-skeletal muscle, we characterized the cellular heterogeneity of endodermal derivatives and inferred the transcriptional events that underlie the specification of endodermal cell types during zebrafish development. We iteratively subclustered 12,592 endodermal cells into 33 clusters and annotated them based on expression of known markers (Fig. 5a, Supplementary Fig. 11a– b). Additionally, we identified the distinct transcription factors expressed by each of these clusters (Supplementary Fig. 11b). The primary source of heterogeneity varied across distinct endodermal tissues. Within the endocrine and exocrine pancreas, clusters primarily reflected distinct cell types. Within the liver, clusters primarily reflected metabolic specialization, such as emergence of a subpopulation of hepatocytes specialized for cholesterol biosynthesis (*msmo1*+) at 84 hpf that persist throughout larval development (Farnsworth, Saunders, and Miller 2020) and until adulthood (Morrison et al. 2022). Within the intestine, clusters were strongly associated with both anterior-posterior position and distinct cell types.

**Figure 5:**
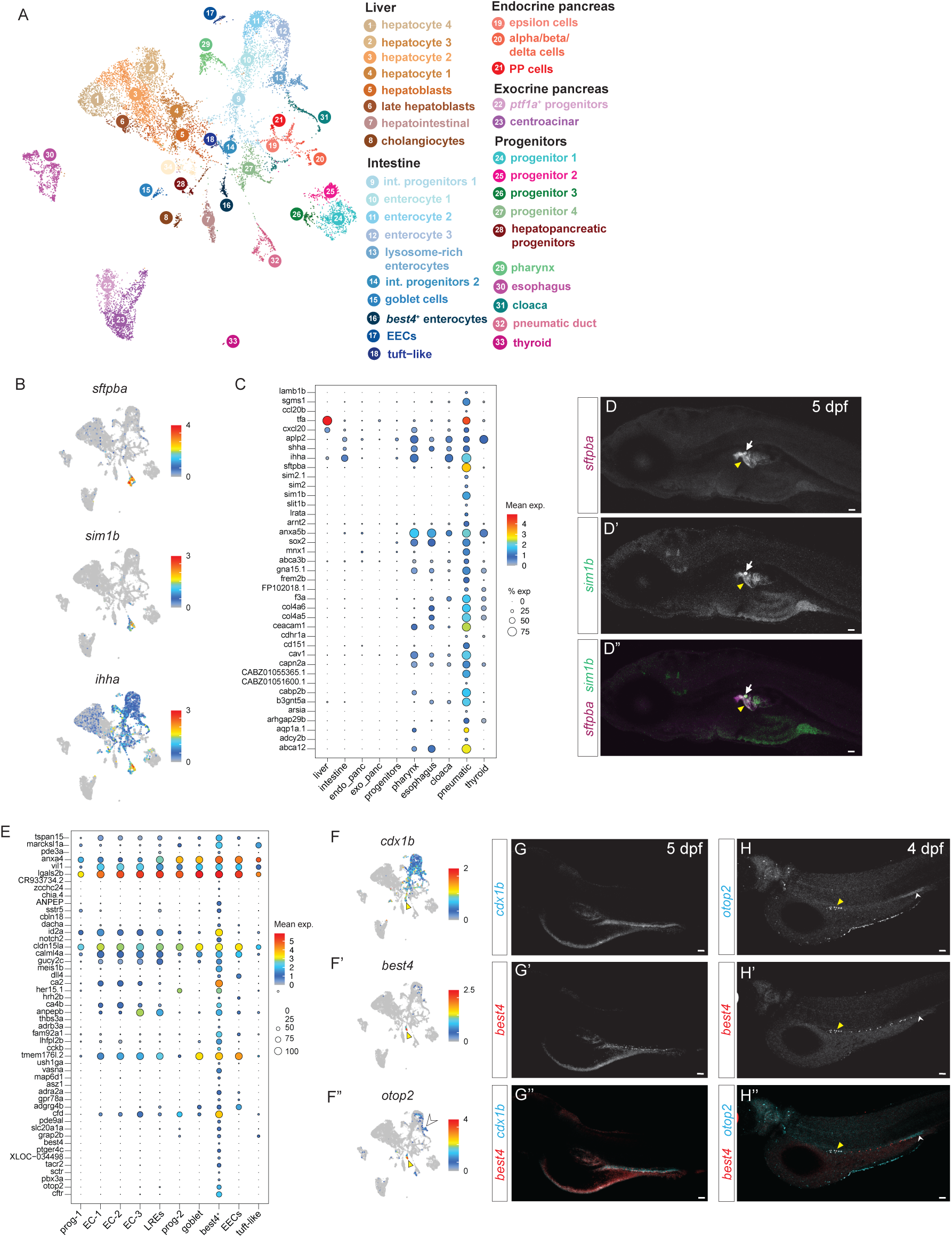
Subclustering of endodermal derivatives enables molecular characterization of human disease associated cell types. (**A**) UMAP projection of 12,592 endodermal cells during zebrafish development, color coded and numbered by cluster. (**B**) Expression of specific (*s*ftpba, *sim1b*) and strongly expressed (*ihha*) pneumatic duct markers visualized on the UMAP projection. Color bar shows expression of each gene. (**C**) Dot plot of top differentially expressed genes (y-axis) within the pneumatic duct compared to other endodermal derivatives (x-axis). (**D–D”**) RNA *in situ* hybridization of two specific markers of the pneumatic duct (*s*ftpba and *sim1b*). Yellow arrowheads indicate staining in the pneumatic duct and inflated posterior swim bladder, white arrows denote staining in the anterior swim bladder bud primordium that inflates at 21 dpf. (**E**) Dot plot showing top differentially expressed markers (y-axis) in *best4*^+^ enterocytes compared to other intestinal cell types. (**F**) Expression of general intestinal marker (*cdx1b*) and two *best4*^+^ enterocyte markers (*best4* and *otop2*) shown on UMAP projection. White arrowhead indicates *otop2* expression in the posterior lysosome-rich enterocyte (LRE) cluster, and yellow arrowhead indicates expression within the *best4*^+^ enterocytes. (**G–H”**) RNA *in situ* hybridization of *best4*^+^ enterocyte marker (*best4*) against (**G–G”**) *cdx1b* and (**H–H”**) *otop2*. Yellow arrowheads indicate *best4* and *otop2* co-expressing cells in the anterior intestine, white arrowheads indicate a posterior patch of *otop2* staining likely corresponding to the posterior LREs as shown in (**F**). Scale bar: 50 µm. **EC**: enterocyte; prog: progenitors; **LREs**: lysosome-rich enterocytes; **EECs**: enteroendocrine cells; **PP**: pancreatic polypeptide

Among endodermal derivatives, the least transcriptionally characterized are those that allow fish to regulate their buoyancy: the swim bladder and the pneumatic duct, which connects the swim bladder to the esophagus. We identified a cluster (C32) that arises at 2–3 dpf which expressed genes that have been previously reported in both the pneumatic duct and swim bladder (*anxa5b*, *hb9/mnx1*, *ihha*, *shha, sox2*) (Fig. 5b–c), but not genes that have been previously reported exclusively in the swim bladder (*acta2*, *elovl1a*, *fgf10*, *has2*, *hprt1l*, *ptch1*, *ptch2*) (Winata et al. 2009; Yin et al. 2011) suggesting that this cluster represents the pneumatic duct. We did not observe a cluster that expressed known swim bladder markers, suggesting that this tissue may not have effectively dissociated during our experiments. Differential gene expression testing revealed that *surfactant protein ba* (*s*ftpba) and the transcription factor, *sim1b* are distinctly expressed in cluster C32 (Fig. 5b–c). RNA *in situ* hybridization confirmed that *s*ftpba and *sim1b* are expressed exclusively in the pneumatic duct and uninflated anterior swim bladder bud (Fig. 5d). While somewhat less specific, pneumatic duct cells also expressed several additional transcription factors (*arnt2*, *sim2*, *sim2.1*) that may be important for its specification. Additionally, consistent with the previously hypothesized evolutionary relationship between the swim bladder and human lung, pneumatic duct cells expressed several genes associated with lung disease, including *ceacam1*, *cd151*, and *abca12* (Fig. 5c). Despite its morphological appreciation for at least a century (Moser 1903), these findings represent (to our knowledge) the first specific molecular markers for the pneumatic duct.

Within the intestine, in addition to three general enterocyte populations, we also detected two non-canonical enterocyte populations (Fig. 5a), including the lysosome-rich enterocytes that are specialized for protein absorption (Fig. 2b). Additionally, we find a zebrafish cell population homologous to human *best4^+^* enterocytes (Fig. 5e–h), a recently described cell type that is potentially reduced in inflamed intestines (Parikh et al. 2019; Smillie et al. 2019). These cells were also very recently identified in 6 dpf zebrafish (Willms et al. 2022), but we first observe them at 3 dpf, when they are in the process of being specified. Similar to their human counterparts, zebrafish *best4+* enterocytes strongly expressed *best4*, *otop2*, *cftr*, carbonic anhydrases (*ca2* and *ca4b* instead of *ca4/7*), and *notch2* (Fig, 5e–f; Fig. 6a–b; Supplementary Fig. 12) (Burclaff et al. 2022; Smillie et al. 2019). Additionally, they strongly express the hormone cholecystokinin (*cckb*) and hormone receptors such as secretin receptor (*sctr*), prostaglandin E receptor 4 (*ptger4c*), adrenergic receptor (*adra2a*), and tachykinin receptor 2 (*tacr2*) (Fig. 5e, Fig. 6b, Supplementary Fig. 12). Via *in situ* hybridization, we found *best4*^+^ cells throughout the intestine, evidenced by overlap with the general intestinal marker *cdx1b* (Fig. 5f, g–g’’)*. best4+* cells in the anterior intestine co-express *otop2* (Fig 5h–h’’, yellow arrowhead), which is also expressed in a posterior patch that is likely the LREs (Fig. 6f’’, white arrowhead). This mirrors recent human single-cell studies that describe *otop2* expression within *best4*+ cells only within particular parts of the intestine, though that location is posterior in humans (the colon) and anterior in zebrafish (Busslinger et al. 2021; Elmentaite et al. 2021).

**Figure 6:**
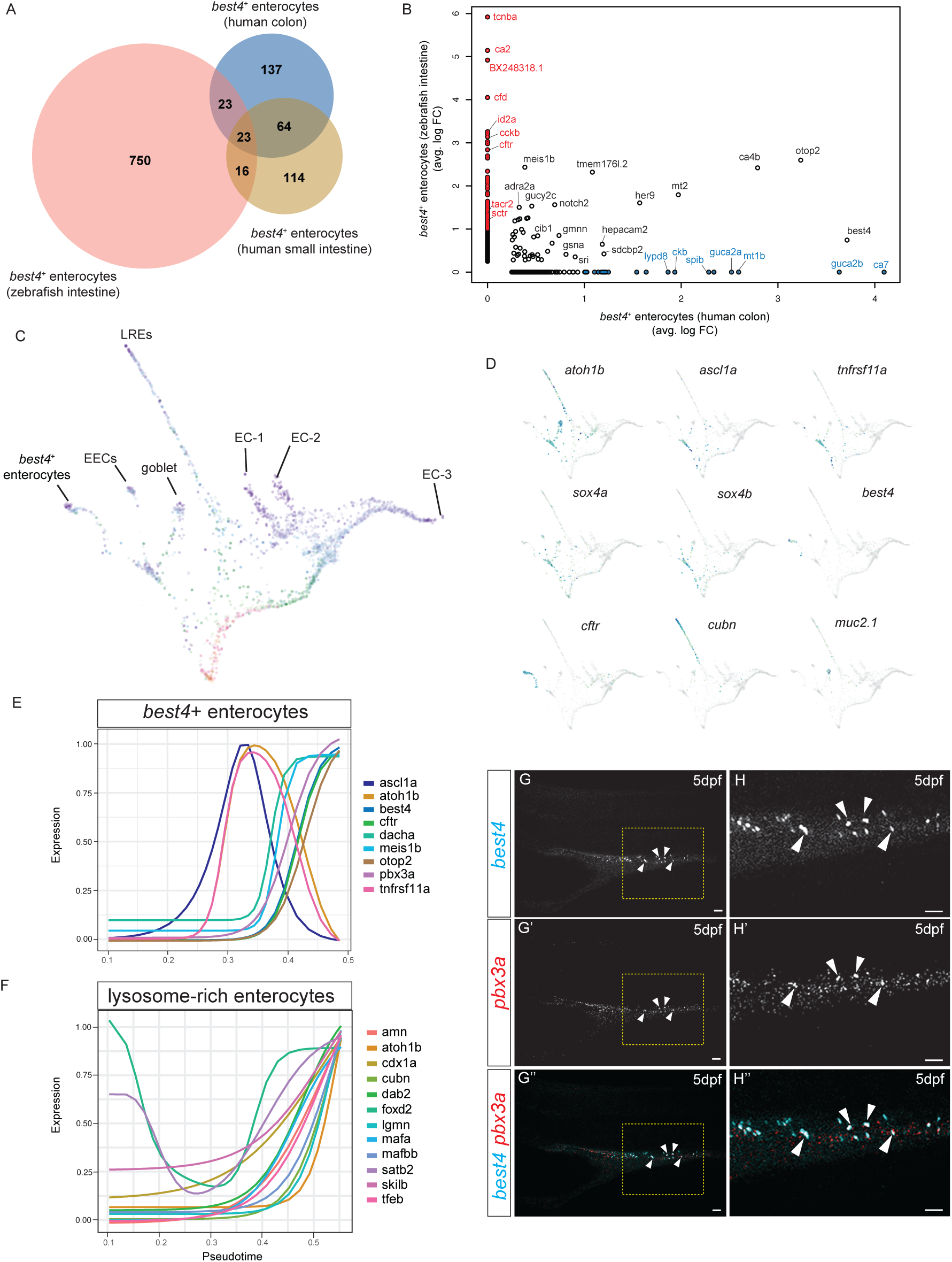
Trajectory analysis of zebrafish intestinal cells reveal candidate regulators for *best4*^+^ enterocyte specification. (**A**) Venn diagram showing number of differentially expressed genes shared between human colonic (Smillie et al., 2019), human small intestinal (Burclaff et al., 2022), and zebrafish *best4*^+^ enterocytes. (**B**) Average log-fold enrichment of genes in *best4*^+^ enterocytes compared to *best4*^+^ enterocyte subtypes in human colon (x-axis, Smillie et al. 2019) and zebrafish intestine (y-axis). Blue: human colon-specific *best4*^+^ enterocyte markers; red: zebrafish-specific *best4*^+^ enterocyte markers; black: shared markers. Comparison of zebrafish *best4*^+^ enterocytes to human small intestinal *best4*^+^ enterocytes shown in Supplementary Figure 12. (**C**) Force-directed layout of a URD-inferred hierarchical tree generated with zebrafish intestinal cells between 14–120 hpf, colored by developmental stage as in Fig. 1A. (**D**) Expression of transcription factors and characteristic markers of individual intestinal cell types visualized on the URD trajectory. (**E, F**) Temporal dynamics of selected genes along the *best4*^+^ enterocyte (E) and posterior LRE (F) trajectory; lines show impulse response fits across pseudotime. Y-axis: scaled expression. (**G–H’’**) RNA *in situ* hybridization of *best4* and candidate transcriptional regulator *pbx3a* in a 5 dpf old zebrafish intestine. (**H–H”**) Higher magnification of yellow boxes from G–G”. Scale bar – 100 µm.

While these non-canonical enterocytes have now been found across multiple organisms, the transcriptional events underlying their specification is unknown. Thus, to establish candidate regulators of these populations, we reconstructed the developmental trajectories of transcriptional changes among intestinal cells using URD and identified transcription factors with dynamic expression along trajectories toward the two non-canonical enterocyte populations (Farrell et al. 2018) (Fig. 6c–f, Supplementary Fig. 13). In progenitors at the branchpoint that separates *best4^+^* cells from other absorptive enterocytes, we observed expression of the RANKL receptor *tnfrsf11a* and the transcription factors *atoh1b* and *ascl1a* (Fig. 6d–e). This is followed by the decline of *ascl1a*, *atoh1b* and *tnfrsf11a* expression, alongside the upregulation of several TFs (including *dacha* and *pbx3a*, whose expression within the intestine is quite specific to *best4+* enterocytes), and lastly expression of cell type specific markers such as *best4*, *otop2*, and *cftr* (Fig. 6e, Supplementary Fig. 13). *In situ* hybridization confirmed that some *best4*+ enterocytes along the mid-intestinal region express *pbx3a* (Fig. 6g–h’’). In other contexts, Pbx3 cooperates with Meis1 to activate transcription (Garcia-Cuellar et al. 2015; Li, Chen, et al. 2016), and we observed that *best4+* cells also express *meis1b* uniquely among intestinal cells (Fig. 5e), and propose that *pbx3a* and *meis1b* may act combinatorially in this cell type as well.

The progenitors that putatively generate LREs were less distinct in their transcriptional state, but we observed an initial decline in expression of the more broadly expressed TFs *foxd2* and *satb2* and upregulation of *cdx1a* and *skilb*. This was followed by upregulation of several TFs (*mafa*, *mafbb*, *tfeb*, *atoh1b*) and genes functionally characteristic of LREs (*cubn*, *dab2*, *amn*) (Fig. 2b, Fig. 6f). Expression of *mafbb* and *tfeb* was unique to LREs among intestinal cells. *tfeb* regulates lysosome biogenesis (Lapierre et al. 2013; Lister et al. 2011)—the most characteristic feature of these cells—suggesting that these other TFs may be involved in either specification of LREs or their functional specialization.

In summary, we catalogued endodermal derivatives during zebrafish development, describing the gene expression program of the molecularly uncharacterized pneumatic duct and multiple non-canonical enterocyte populations in zebrafish, including *best4*+ enterocytes. We established the molecular similarity of zebrafish and human *best4*+ enterocytes and used trajectory approaches to identify putative progenitor populations and candidate developmental regulators that may be important for the specification or differentiation of these enterocyte populations.

## Discussion

Understanding how gene expression changes drive specification and differentiation of distinct cell types during animal development benefits massively from a complete understanding of the cast of players: which molecular cell types/states exist, and which genes are expressed by each. In this study, we mapped gene expression landscapes with high temporal resolution across 489,686 cells during the first 120 hours of zebrafish development. Analysis of these data revealed global gene expression programs, characterized the persistence of transcriptional states during development, identified undescribed or rare cell types/subtypes, and enabled molecular characterization of tissues with few known marker genes. Moreover, we generated testable hypotheses about crucial regulators of cellular function and cell specification. We anticipate that additional focused re-analyses of these data by other investigators will lead to further discoveries.

One application of large-scale, whole-animal timecourse single-cell data is that it enables comparison between different cell types, tissues, and developmental times at the whole-transcriptome level. This enabled us to build a catalog of gene expression program that are reused by multiple cell types during development. Additionally, this allowed us to assay whether developmental transcriptional states are found only briefly or can be found over many developmental stages. We identified that most transcriptional states are found for 12–36 hours (based on the threshold for transcriptional similarity that we used). Additionally, this analysis helped separate stem cell populations into groups that have a stable transcriptional state and groups whose transcriptomes change during development (such as progenitor cells in the ventral nerve tube or mouse epiblast, which produce different descendants at different times). The stem cell populations which we identify as having a “long-term” transcriptional state are likely to remain transcriptionally consistent at the whole-transcriptome level. We find that this applies to several classic stem cell populations usually identified by a few markers, including radial glia, satellite cells, and several populations within the eye. Moreover, our analysis captured a few additional populations that have not been studied as stem cells that occupy a “long-term” dividing state. These candidates include *oit3*^+^ lymphatic endothelia, *pah*^+^/*hpdb*^+^ dermal fibroblasts, myoseptal fibroblasts, and cephalic/opercular muscles. Further experimental investigation may establish these states as true progenitor populations.

In addition to such global insights into zebrafish development, focused analyses in this work revealed the gene expression profiles of cellular subtypes that have not been well characterized molecularly, including the pneumatic duct and intestinal smooth muscle layers. These provide the molecular handles required for dissecting the development and function of these tissues. For instance, we identify genes that are specific to the pneumatic duct, including transcription factors that may regulate its development (*sim1b, sim2*) and Sftpb surfactant protein, which prevents terminal airway collapse while breathing in human lungs (Stahlman et al. 2000; Han and Mallampalli 2015). The pneumatic duct expresses other genes associated with pulmonary surfactant metabolism dysfunction, such as *abca3* (Ban et al. 2007), which suggests that the swim bladder may be a valuable model for understanding and developing therapeutics to treat these disorders. This motivates testing the function of other specific markers of this tissue; for instance, tissue-specific models of *abca12* may reveal a role in lung function and surfactant metabolism, in addition to its known role in maintaining epidermal barrier function (Zuo et al. 2008). Similarly, though the developmental signaling regimes that generate distinct intestinal smooth muscle layers are known (Huycke et al. 2019), this study identified transcriptional differences that are potentially downstream, including several candidate TFs. Interestingly, many of the TFs restricted to circular smooth muscle (*foxq1a*, *foxq1b*, *tcf21*) inhibit SMC differentiation in mammals by opposing the activities of the Foxf and Myocardin pathways (Hoggatt et al. 2013) which we observe in both intestinal SMC types. This presents the intriguing possibility that a key aspect of circular smooth muscle differentiation may involve the inhibition of alternative SMC fates, including longitudinal muscle. Our identification of distinct markers of the individual intestinal SMC layers will enable their genetic manipulation to test their functional contributions, developmental origins, and the regulation underlying their specification.

For some cell types, focused analysis revealed previously unappreciated heterogeneity that has potential functional implications. For instance, although pericyte morphological heterogeneity is well appreciated (Grant et al., 2019), pericyte molecular heterogeneity remains an area of investigation, partially due to the former dearth of pericyte specific markers, which was recently resolved (Shih et al. 2021). This study reveals that zebrafish possess multiple transcriptionally distinct pericyte subpopulations. Notably, at least some of the subtype-specific genes that we observe are expressed in pericytes in other animals (for instance, *adma* in rat pericytes (Kis et al. 2002) and *epas1a* and *esama* in mouse pericytes (Muhl et al. 2020)). In systems where transcriptional differences have been identified among pericytes, they are often accompanied by spatial restriction; for example, mouse lung and brain pericytes differ in expression of several transmembrane transporters (Vanlandewijck et al. 2018). Similarly, this study observes that the pericyte-2 subtype is associated with particular vessels in the hindbrain, suggesting that zebrafish pericytes may also exhibit tissue-dependent transcriptional differences. Moreover, the genes expressed in different pericyte populations observed in this study may indicate distinct functions for these subtypes that will need to be tested using reverse genetic approaches. For example, we observe that only a subpopulation of zebrafish pericytes express the peptide hormone adrenomedullin (*adma*), which is produced by rat cerebral pericytes, where it triggers vasodilation of the vessels they surround (Kis et al. 2002). Similarly, since pericyte-2 (*epas1a*+) pericytes were often observed in pairs and sometimes at some distance from the vessel, this suggests that pericyte-2 subtype gene may be associated with particular morphological or migratory characteristics. Future long-term imaging assays will be required to determine whether these distinct transcriptional profiles represent persistent pericyte identities or transient cell states that pericytes in certain anatomical locations enter and exit.

This study also reconstructed trajectories that describe the development of recently discovered and potentially disease-related cell types, such as the *best4*^+^ enterocytes (Parikh et al. 2019; Elmentaite et al. 2021). Decreased numbers of *best4*+ enterocytes in patients with ulcerative colitis and disruption of cGMP signaling in colorectal cancer (Parikh et al. 2019; Pattison et al. 2020; Smillie et al. 2019) have suggested a potential importance for *best4*+ enterocytes in human disease. *Best4*+ enterocytes are not found in rodents (Ikpa et al. 2016; Nowotschin et al. 2019; Pijuan-Sala et al. 2019), suggesting that zebrafish represent an excellent opportunity to study the function and development of these cells. Comparison between human and zebrafish *best4*+ enterocytes revealed important transcriptional similarities that indicate functional homologies. Human *best4*+ enterocytes respond to luminal pH (Parikh et al 2019) and zebrafish share the *best4* and *otop2* ion channels that are proposed to mediate this response. Additionally, human and zebrafish *best4*+ enterocytes share expression of *cftr* and carbonic anhydrases which may regulate ion or fluid homeostasis in the gut. Lastly, shared expression of *adra2a* between both human and zebrafish *best4^+^* enterocytes (Supplementary Fig. 12) may indicate a role in intestinal motility in both organisms (Scheibner et al. 2002). Human *best4*+ enterocytes are the source of the critical hormone Uroguanylin/Guca2b (a guanylate-cyclase agonist peptide that increases cGMP levels to regulate satiety and intestinal tone) (Valentino et al. 2011). Although zebrafish guts do not express an ortholog of Uroguanylin, zebrafish best4+ enterocytes instead produce the intestinal hormone cholecystokinin which also regulates satiety, intestinal pH, and intestinal motility in part by activating cGMP production (Sindic 2013; Konturek, Konturek, and Domschke 1995; Zeng et al. 2020). This suggests that a conserved function between human and zebrafish best4+ enterocytes may be to activate cGMP production in response to pH changes or other signals, albeit via different mechanisms. Lastly, human *best4*+ enterocytes in different regions of the intestine exhibit distinct gene expression profiles (Burclaff et al. 2022; Parikh et al. 2019; Smillie et al. 2019). Low cell numbers prevented sub-clustering the *best4*+ enterocytes in this study, but *in situ* hybridization demonstrated that zebrafish also exhibit regionalization of this population, since *otop2* co-expression is limited to particular regions of the intestine in both zebrafish and humans. Understanding the biology of these cells—both their development and function—would be crucial to enabling therapeutic approaches that manipulate or target them. Thus, in this study, we identify the molecular characteristics of a putative progenitor state that gives rise to these cells (*atoh1b/ascl1a/tnfrsf11a*+), identify candidate developmental signals and regulators that may be important for their specification (e.g. Notch2, BMP, *dacha*, *pbx3a*, *meis1b*, *tox2*), and identify cell-type specific markers of these cells. Altogether, these lay the groundwork for experiments and genetic tools to manipulate these cells to understand their development and function in intestinal homeostasis and cell-cell communication with neighboring cell types.

As exemplified in this study, single-cell genomics approaches have seen wide adoption among developmental biologists, and these data join a few complementary single-cell RNAseq datasets focused on zebrafish development. While all profile whole animals, each study has focused on analyzing different tissues or developmental processes, such as the early embryo and axial mesoderm (Farrell et al. 2018), early embryo and pharyngeal arch (Wagner et al. 2018), the intestine and non-skeletal muscle (this work), liver and notochord (Farnsworth, Saunders, and Miller 2020), or parachordal cartilage and cranial ganglia (Saunders et al. 2022). This highlights the richness of the data and the potential for its reuse and reanalysis to make additional discoveries. It also underscores the need to make these data accessible and easy to work with. To this end, we created Daniocell to enable researchers to browse our data rapidly to address simple questions.

We anticipate that these data can augment other large-scale efforts to build models or understand mechanisms of development. For instance, we envision that label transfer approaches (Stuart et al. 2019; Lotfollahi et al. 2022) used with our annotations will accelerate the time-consuming annotation step of future single-cell RNAseq work in zebrafish—at least to some degree; for this purpose, we encourage others to submit improvements to the annotations through Daniocell. Integration with data generated from different transgenic lines or via CRISPR barcoding techniques (McKenna et al. 2016; Spanjaard et al. 2018) can ascribe lineage information to each cell population and help understand their developmental origins. These data also provide a framework for unraveling the gene regulatory network underlying vertebrate development, especially when combined with high-throughput single-cell chromatin accessibility assays and computational gene regulatory network inference approaches (Aibar et al. 2017; Kamimoto et al. 2023). Thankfully, increasingly high-throughput CRISPR screening techniques mean that approaches to functionally test the predictions that would emerge from those efforts are also accelerating (Parvez et al. 2021). The greatest gains from these approaches will be realized as cooperation increases within the zebrafish community to collectively integrate the results generated across many labs using different techniques, stages, and transgenic and mutant lines to make them more broadly useable.

## Materials and Methods

### Zebrafish Husbandry

This study includes the use of live zebrafish vertebrate embryos. Animals were handled according to National Institutes of Health (NIH) guidelines. Some zebrafish work was performed at the facilities of Harvard University, Faculty of Arts & Sciences (HU/FAS) under protocol 25-08. The HU/FAS Institutional Animal Care and Use program maintains full AAALAC accreditation, is assured with OLAW (A3593-01) and is currently registered with the US Department of Agriculture (USDA). Additional zebrafish work was performed at the *Eunice Kenndy Shriver* National Institute of Child Health and Human Development (NICHD) Shared Zebrafish Facility, under animal protocol 20-001. The NICHD Animal Care and Use program also maintains full AAALAC accreditation, is assured with OLAW (D16-00602) and is currently registered with the US Department of Agriculture (USDA). At the developmental stages profiled in this study, zebrafish sex is not yet determined, so sex was not considered a biological variable in this study.

### Dissociation of animals into cell suspensions

Fertilized eggs were collected from wild-type (Tupfel long fin/AB) in-crosses and then cultured in blue water (5 mM NaCl, 0.17 mM KCl, 0.33 mM CaCl2, 0.33 mM MgSO4, 0.1% methylene blue) at 28°C until they reached the desired stage. Per sample, 10–12 embryos or larvae were dechorionated using forceps in calcium-free Ringer’s solution (116 mM NaCl, 2.6 mM KCl, 5 mM HEPES, pH 7.0, 500 mL, sterile) with MESAB (400 ug/mL) and left to sit 5–10 minutes at 28°C to become anesthetized. Protease solution was mixed fresh each day (0.25% trypsin, 1 mM EDTA, pH 8.0, in 1xPBS, sterile; Sigma-Aldrich T4549). Per sample, 1.2 mL of protease solution was added to a well in a 24-well plate. This plate was placed at 28°C to equilibrate temperature. Meanwhile, collagenase P solution (100 mg/ ml Collagenase P in Hank’s Balanced Salt Solution; Sigma-Aldrich H9269 and 11213865001) was thawed on ice. Each sample of embryos/larvae were transferred to a 1.5mL Eppendorf tube, and the volume of Ringer’s solution was reduced to 100 uL. The animals were de-yolked and abraded by pipetting up and down 5 times with a P200 set to 80 uL. Animals were rinsed by adding 1 mL of fresh Ringer’s solution, allowing the animals to settle to the bottom, and removing all but 100 uL of the Ringer’s solution. The animals (including the 100 uL of Ringer’s) were then transferred into the 24-well plate with protease solution, 30 uL of Collagenase P solution was added per well, and the plate was swirled to mix (final volume = 1.33 mL per well). The plate was incubated at 28°C, monitoring dissociation with a microscope, pipetting up and down with a P1000 20 times slowly every 5 minutes. After 10 minutes, each sample was passed once through a P200 by pipetting from one well into another. Digestion was stopped after 20–25 minutes by adding 270 uL of 6X STOP solution (6X, 30% calf serum, 6 mM CaCl2, in 1 x PBS, sterile) to each well and swirling gently to mix (final volume per well now 1.5 mL). In many cases, some small chunks of tissue remained, but attempting to achieve complete digestion usually reduced cell viability significantly. Suspensions were filtered through a 40 uM cell filter into 1.5 mL Eppendorf tube to remove any remaining tissue chunks. The tube was flicked vigorously 20 times.

### MULTI-seq barcoding of cell suspensions

Cells were spun down at 300 x g for 3 minutes at 4°C. Supernatant was removed (leaving ∼ 100 ul) and cells were resuspended in 1 mL chilled DMEM/F12 (0% BSA; Gibco 12500062). Cells were washed by spinning at 300 x g for 3 minutes at 4°C, removing supernatant, and resuspending cells in 1 mL chilled DMEM/F12 (0% BSA). During the spin, 11 uL of anchor solution was added to each well of a round-bottom plate (200 nM anchor oligo, 200 nM barcode oligo in DMEM/F12 with 0% BSA; anchor oligo was a gift from Chris McGinnis and the Zev Gartner lab, now available from Sigma-Aldrich; barcodes ordered from IDT). Cell suspension was distributed into 10-12 wells (100 uL cells per well), mixed with 5-10 gentle pipettings, then incubated on ice 5 minutes. To lock MULTI-seq barcodes in place, 11 uL of co-anchor oligonucleotide solution (200 nM co-anchor oligo in DMEM/F12 with 0% BSA; co-anchor oligo also a gift from Chris McGinnis and the Zev Gartner lab, now available from Sigma-Aldrich) was added to each well, pipetted gently to mix, and incubated on ice for 5 minutes. Labeling was halted with BSA by adding 50 uL of DMEM/F12 (4% BSA) and mixing gently. To remove excess labeling reagents, cells were washed twice by spinning cells down at 300 x g for 3 minutes at 4°C, removing supernatant, and resuspending in 200 uL of DMEM/F12 (1% BSA). After the final wash, samples were pooled in a 2 mL Eppendorf, washed by spinning at 250 x g for 4 minutes, removing the supernatant, and resuspending in 1 mL of DMEM/F12 with 1% BSA, spinning again at 250 x g for 4 minutes, and then resuspending in 300 uL of DMEM/F12 with 1% BSA. Cells were filtered through a 40 uM FlowMi cell filter (Bel-Art H13680-0040) into a clean tube.

### Collection of single-cell transcriptomes

Droplet emulsions of single cells were generated using the 10X Genomics Chromium controller with Single Cell 3’ v3.1 consumable reagents, according to the manufacturer’s instructions. In brief, single-cell suspensions were stained with acridine orange / propidium iodide (11 uL of cells + 1 uL of Logos F23001 Acridine Orange/Propidium Iodide Stain) and then examined and quantified on a Logos Luna FL automated cell counter. Samples with sufficient concentration, low multiplet rate (<5% to proceed, <3.6% on average), and high viability (>85% to proceed, >95% on average) were then diluted with Ringer’s to load into the instrument. Cells had been barcoded with MULTI-seq hash oligos to enable overloading of the instrument, which would then return of a larger number of transcriptomes with an increased rate of doublets that could then be removed computationally based on identification of multiple hash barcodes associated with that cell barcode. Samples were loaded into two channels at normal (9,600 cells loaded, targeting 6,000 cells recovered) and high (34,000 cells loaded, targeting 21,250 cells recovered) concentrations. Downstream reactions were performed in the Biorad C1000 Touch Thermal cyclers. 10–12 cycles were used for cDNA amplification, and the result was inspected using Agilent High Sensitivity DNA Kits on the Agilent Bioanalyzer 2100. 8–13 cycles were then used for sample index PCR, based on the concentration of amplified cDNA in the previous step. Final libraries were evaluated using Agilent High Sensitivity DNA Kits on the Agilent Bioanalyzer 2100 and quantitated using a ThermoFisher Qubit 4 with dsDNA High Sensitivity (HS) reagents. Average fragment length from the Bioanalyzer and concentration from the Qubit were used to pool libraries in equimolar concentrations. Libraries were sequenced across several sequencing runs. Some samples (including samples TC1-24, TC1-48, TC1-72, TC1-96, TC2-36, TC2-60, TC2-84, and TC3-48) were checked in three separate runs on a Nextseq 500 System (Illumina) with High Output 75 cycle kits, with 28 bases for Read 1, 8 bases for Index 1, and 56 bases for Read 2. Most sequencing was performed in three separate runs on a NovaSeq 6000 Sequencing System (Illumina), using S4 full flowcells, with 28 bases for Read 1, 8 bases for Index 1, and 91 bases for Read 2. PhiX control library was spiked in at 1%. Libraries were briefly analyzed and re-pooled after the second sequencing run to try to achieve similar reads/cell across the entire data. Reads from all runs above were used in downstream analysis. Most sequencing runs contained mixtures of libraries to measure gene expression and MULTI-seq barcodes.

### Alignment of sequencing data

Alignment of sequencing reads and processing into a digital gene expression matrix was performed using Cell Ranger version 4.0.0, including the aligner STAR version 2.5.1b, with standard parameters. The-expect-cells parameter was set to 6,000–21,250 based on the number of cells loaded per sample. The data was aligned against GRCz11 release 99 (January 2020) using the Lawson Lab Zebrafish Transcriptome Annotation version 4.3.2, published in Lawson et al. eLife 2020, available from https://www.umassmed.edu/lawson-lab/reagents/zebrafish-transcriptome/. 320 entries annotated as pseudogenes by Ensembl were removed from the reference.

### Removal of MULTI-seq doublets

UMI count tables of MULTI-seq cell hashing barcodes were generated using the deMULTIplex package (available https://github.com/chris-mcginnis-ucsf/MULTI-seq/). Reads were input from FASTQ and preprocessed (deMULTIplex::MULTIseq.preProcess, cell=c(1,16), umi=c(17,28), tag=c(1,8)), aligned against the barcode sequences used in each experiment and then deduplicated into a UMI counts table (deMULTIplex::MULTIseq.align). Visual inspection on a tSNE projection (deMULTIplex::barTSNE) was used to confirm that the run had been successful and cells fell into clearly defined barcode classes.

In order to remove doublets that resulted from overloading 10X channels in barcoded samples, MULTI-seq encoded cell hashes were used to remove resultant doublets. Briefly, two calculations were used — classification based on Seurat’s hash tag oligo demultiplexing functions, and a classification based on signal-to-noise ratio. For classification by Seurat, a Seurat object was created using the MULTI-seq UMI counts matrix, normalized (Seurat::NormalizeData, assay=“MS”, normalization.method = “CLR”), and classified (Seurat::HTODemux, assay=“MS”, positive.quantile=0.9999). For classification based on signal-to-noise, cells were called as ‘negative’ if they had <20 UMIs aligned to a single barcode. To be considered a ‘singlet,’ required that the signal-to-noise ratio was ≥5, where ‘signal’ was the number of UMIs assigned to the barcode with the most UMIs and ‘noise’ was the number of UMIs assigned to the barcode with the second most UMIs. Cells with signal-to-noise ratio <5 were classified as ‘doublets.’ Cells were removed for lacking MULTI-seq cell hash information if they were called as ‘negative’ in both approaches. Cells were removed for being doublets if they were scored as a ‘doublet’ by *either* approach.

### Normalization and quality control

Cells that were scored as singlets based on MULTI-seq cell hashing were then processed and analyzed using Seurat version 3.1.5 and R version 3.6.3. First, cells were scored (Seurat::PercentageFeatureSet) for their mitochondrial gene expression (using all genes beginning *mt-*) and ribosomal gene expression (using the genes: *rpl18a, rps16, rplp2l, rps13, rps17, rpl34, rpl13, rplp0, rpl36a, rpl12, rpl7a, rpl19, rps2, rps15a, rpl3, rpl27, rpl23, rps11, rps27a, rpl5b, rplp2, rps26l, rps10, rpl5a, rps28, rps8a, rpl7, rpl37, rpl24, rpl9, rps3a, rps6, rpl8, rpl31, rpl18, rps27.2, rps19, rps9, rpl28, rps7, rpl7l1, rps29, rpl6, rps8b, rpl10a, rpl13a, rpl39, rpl26, rps24, rps3, rpl4, rpl35a, rpl38, rplp1, rps27.1, rpl15, rps18, rpl30, rpl11, rpl14, rps5, rps21, rpl10, rps26, rps12, rpl35, rpl17, rpl23a, rps14, rpl29, rps15, rpl22, rps23, rps25, rpl21, rpl22l1, rpl36, rpl32, rps27l*).

Cells were removed with either a low number of detected features (≤200 genes detected) or abnormally high number of detected features (top 0.5%), or with abnormally high mitochondrial content (≥10%). They were then log-normalized (Seurat::NormalizeData, normalization.method = “LogNormalize”, scale.factor = 10000) and scaled, regressing against mitochondrial and ribosomal gene expression (Seurat::ScaleData, vars.to.regress = c(“percent.mt”, “percent.ribo”)).

### Remapping and integration of 2018 Drop-seq data

For earlier stage cells in this analysis, previously published single-cell transcriptomes from wild-type TL/AB zebrafish covering 3.3–12 hours post-fertilization from Farrell et. al 2018 were integrated with the newly generated data. First, the previous data was re-aligned to the reference used in this study and processed using Drop-seq Tools version 1.12 and its included copy of Picard Tools. Since the final step of the Drop-seq processing pipeline corrects cell barcodes to account for oligonucleotide synthesis errors that occur during the manufacture of the Dropseq beads, we started from the BAM files generated in the 2018 study, where cell barcodes had already been corrected, in order to maintain consistency with the original study. Picard Tools SamToFastq was used to create a FASTQ file from the previous BAM file, which was then used as input to STAR version 2.5.4a with the same reference that had been used with CellRanger. Picard Tools RevertSam was used to remove the previous alignment from the 2018 BAM, then both the 2018 BAM and output of STAR were sorted into queryname order using Picard Tools SortSam (SO=queryname). The remaining steps were standard application of Dropseq Tools. The two BAM files were merged with PicardTools MergeBamAlignment, tagged with gene exon information using Dropseq Tools TagReadWithGeneExon and a digital gene expression matrix was produced using Dropseq Tools DigitalExpression with NUM_CORE_BARCODES = 12000. The digital gene expression matrices were then combined and trimmed to match exactly the cells included in the original 2018 study.

Using Seurat version 4.1.0, separate Seurat objects were created for the remapped 2018 Dropseq dataset (Farrell et al. 2018) and newly generated 10X dataset in this study. These objects were combined using the SeuratObject::merge command. For visualization, we identified the top 2000 variable genes (Seurat::FindVariableFeatures, selection.method = “vst”, nfeatures = 2000), performed PCA (Seurat::RunPCA), and identified significant PCs (Seurat::JackStraw, dims=100). A Uniform Manifold Approximation and Projection (UMAP) was calculated using 50 nearest neighbors and the 30 most significant PCs (Seurat::RunUMAP, n.neighbors = 50). For clustering, we used an iterative approach. First a broad clustering (“top clusters”) was generated by identifying the top 1500 variable genes (Seurat::FindVariableFeatures, selection.method = “vst”), performing PCA, and clustering using the Leiden approach on the top 30 PCs (Seurat::FindClusters, algorithm = 4, resolution = 0.1, n.start = 50, random.seed = 17), resulting in 25 clusters. Within each top cluster, sub-clusters were determined using a similar approach, by calculating the top 2000 variable genes, performing PCA, identifying the significant PCs, and performing Leiden clustering at multiple resolutions (Seurat::FindClusters, algorithm=4, resolution = c(0.75, 1, 2, and 3), n.start = 50, random.seed = 17). Markers were calculated for each subclustering and roughly annotated subclusters. The different resolutions were assessed manually, and a resolution was chosen based on which clustering seemed the most biologically relevant. Most often, clusters obtained at resolution 2 and 3 were found to best capture cell type differences within each subclustering, but within some less-complex tissues (*e.g.* the primordial germ cells), lower resolutions were more appropriate. In order to make the “top clusters” biologically relevant and represent individual tissues within the fish (for instance, “endoderm”), some subclusters were manually reassigned to different top clusters based on their cell type annotations, and some top clusters were manually split. The subclustering procedure was repeated, and final subcluster resolutions were chosen again based on biological relevance. Additional manual curation was performed by manually splitting some subclusters that represented more than one cell type based on their expressed genes and/or prior knowledge from the literature. Additionally, subclusters without sufficient differentially expressed genes were combined in order to ensure that each subcluster was sufficiently distinct: marker genes for individual clusters were identified using ROC and Wilcoxon Rank Sum Tests using the command Seurat::FindMarkers(test.use = “roc”/”wilcox”, min.pct = 0.25, logfc.threshold = 0.25) and subclusters without at least 3 differentially expressed genes were merged. While heavily manually curated, our overall goal was to represent the molecular heterogeneity of cell types recovered in our data while also representing the known biology of zebrafish development to the best of our ability. This final clustering comprised 19 tissue subsets (top clusters), which contained a total of 521 subclusters. The final clusters were annotated based on previously described expression patterns from published RNA *in situ* hybridizations, published single-cell RNAseq studies, public repositories such as ZFIN, and elsewhere. Annotations are all compiled in Table S1 with a nested hierarchy containing information about germ layer/tissue/organs, broader tissue category, and finally the cell type.

### Identifying short-term and long-term transcriptional states during development

To identify groups of transcriptionally similar cells, we used an ε-neighborhood approach. Euclidean distances in gene expression space were calculated between cells in each of the 19 tissue subsets using the stats::dist function. Genes used were the union of the highly variable genes calculated on each of the 19 tissue subsets. In order to find an optimal ε, we assessed neighbors identified for each cell using a range of different ε neighborhood sizes (ε = 20, 25, 30, 31, 32, 33, 34, 35, 36, 37, 38, 39, 40, 41, 42, 43, 44, 45, 50, 55, 60), and chose the smallest ε that allowed most cells in the data to have ε-neighbors. We observed that at least 70% of cells within each of the 19 subsets had neighbors within an ε size of 35.

Each cell was assigned a cell cycle score based on expression of genes representative of G1/S, G2/M, and G0 phases of the cell cycle using the Seurat::CellCycleScoring function. Genes used to assign cell cycle scores to cells in our dataset were as follows: G1/S – *mcm5*, *pcna*, *tyms*, *mcm7*, *mcm4*, *rrm1*, *ung1*, *gins2*, *mcm6*, *cdca7*, *dtl*, *prim1*, *uhrf1*, *cenpu*, *gmnn, hells*, *ccne2*, *cdc6*, *rfc2*, *polr1b*, *nasp*, *rad51ap1*, *wdr76*, *slbp*, *ubr7*, *pold3*, *msh2*, *atad2*, *rad51*, *rrm2*, *cdc45*, *exo1*, *tipin*, *dscc1*, *blm*, *casbap2*, *usp1*, *clspn*, *pola1, chaf1b*, *mrpl36*, *e2f8*; G2M – *cdk1*, *ube2c*, *birc5*, *top2a*, *tpx2*, *cks2*, *nuf2*, *mki67*, *tacc3*, *cenpf*, *smc4*, *ckap4*, *kif11*, *cdca3*, *hmgb2*, *ndc80*, *cks1b*, *tmpo*, *pimreg*, *ccnb2*, *ckap2l*, *ckap2*, *aurkb*, *bub1*, *anp32e*, *tubb4b*, *gtse1*, *kif20b*, *hjurp*, *jpt1*, *cdc20*, *ttk*, *cdc25c*, *kif2c*, *rangap1*, *ncapd2*, *dlgap5*, *cdca8*, *cdca2*, *ect2*, *kif23*, *hmmr*, *aurka*, *psrc1*, *anln*, *lbr*, *ckap5*, *cenpe*, *ctcf*, *nek2*, *g2e3*, *gas2l3, cbx5*, *cenpa.* Cells were then categorized as ‘cycling’ (based on either G1/S or G2/M scores above 0) or ‘non-cycling’ (based on G1/S and G2/M scores both below 0).

Using an ε neighborhood size of 35, for each cell, we identified the ε-neighbors, computed the absolute value of the difference in developmental stage between the cell and each of its neighbors, and then took the mean as that cell’s “stage difference.” The “stage difference” identifies whether for a given cell, the transcriptional similar cells are generally very similar in stage (in which case this value will be small), or transcriptionally similar cells span a wide range of developmental cells (in which case this value will be large). We categorized cells based on “stage difference” into different groups (“<24hr”, “24–36hr”, “36–48hr”, and “≥48hr”), and considered cells with a “stage difference” of ≥36 hours as “long-term” states.

For plots of long-term cycling states in Fig. 1F, clusters with at least 10% of cells in a long-term cycling state were considered. For these states, we identified their ε-neighbors and the clusters to which the neighbors belonged. Long-term cycling states that exhibited ≥99% overlap (i.e. that identified 99% of another cluster as neighbors) were considered similar transcriptional states and were combined. For the neighbors of the resultant long-term cycling states, only those clusters were considered in which at least 50% of cells were neighbors of the long-term state. Clusters associated with long-term cycling states and their neighbors were then aligned with our annotations. Clusters that corresponded to doublets or technical artefacts were also removed from this analysis. The bar represents the shortest time range that encompassed 80% of the “long-term” cells and their transcriptionally similar ε-neighbors.

### Gene module identification

Identification of gene modules is sensitive to the noise inherent in scRNA-seq data. To ameliorate this, we used a form of imputation/smoothing based on *k*-nearest neighbor networks (Wagner, Yan, and Yanai 2018). Briefly, PCA was conducted on the data and the 5 nearest neighbors were identified for each cell based on the top 30 PCs using the command knn_smoothing (k=5, d=30, seed=42) from the R implementation of the Wagner, Yan, and Yanai approach.

Due to limitations on the addressable size of a matrix for the clustering package chosen, we focused on a subset of genes and downsampled the cells used as input. Genes were limited to those that were expressed in 0.1% – 75% of cells in the data to exclude genes who were too lowly expressed to produce meaningful results and to focus on genes that exhibited cell-type specificity by excluding genes that were mostly ubiquitous. To maximize retention of cellular complexity, downsampling was performed to focus on eliminating cells from overrepresented populations while preserving rare cell types and changes over time. Each cluster was divided according to its major “stage groups” (3–4 hpf, 5–6 hpf, 7–9 hpf, 10–12 hpf, 14–21 hpf, 24–34 hpf, 36–46 hpf, 48–58 hpf, 60–70 hpf, 72–82 hpf, 84–94 hpf, 96–106 hpf, 108–118 hpf, 120 hpf). Per cluster–stage-group (i.e. the cells defined by the intersection of cluster and stage group), 50 cells or 20% (whichever was larger) was retained.

Fuzzy c-means clustering was then performed using the R package Mfuzz (Kumar and M 2007). Briefly, data was standardized (Mfuzz::standardise) and then clustered with fuzziness parameter 1.04 (empirically determined) and 200 modules requested (Mfuzz::mfuzz, c = 200, m = 1.04). These were then filtered for technical quality. First, modules that were extremely similar (member gene loadings had correlation >0.95) were combined by summing their member gene loadings (eliminating 48 of 200 modules). Second, intra-cluster variation in expression patterns was reduced by limiting gene memberships to core members (“α-core members” in Mfuzz parlance) that contribute the most strongly to the overall ‘expression’ of a module—gene loadings <0.2 were converted to 0. Third, any modules that had fewer than 5 core member genes were eliminated (eliminating 5 of 152 remaining modules). Finally, new cell embeddings were generated by multiplying the new gene loading matrix against the original expression data (new.cell.embedding <-Matrix::t(data.unlogged) %*% membership.adjusted).

The resultant 147 modules were analyzed for their expression across tissues and how completely individual genes within these modules were shared across tissues. Gene modules were then annotated to identify the functional roles of their constituent genes based on literature and information from public repositories such as ZFIN. Genes from each module were grouped and analyzed together based on their previously reported functions or based on the family that they belonged to. For example, in GEP-193 (Supplementary Table 2), two groups of genes were identified: one associated to Megalin-mediated endocytosis and the other represented a family of SLC transporters. Gene modules that (i) represented outliers, (ii) consisted of member genes expressed only within one tissue subset, and (iii) and contained member genes that were mitochondrial or ribosomal were excluded from our downstream analysis eliminating 57 of the resultant 147 modules.

Gene modules were calculated using all cells as well as cells constituting each tissue type. To compare the gene modules recovered between the global dataset versus the tissue groups, we calculated correlation and cosine similarity between all modules that were calculated either globally or on a particular tissue. Modules with a correlation of at least 0.25 were sorted and clustered using the functions hclust(method = “ward.D2”) and cutree(h = 0.75). All modules calculated on individual tissues were either: (1) highly correlated with a gene module calculated on the global data set, or (2) not recovered in the global analysis, but also not shared with another tissue in the dataset.

### Embryo pretreatment and fixation

Zebrafish larvae (3–5 dpf) were collected and fixed in 4% paraformaldehyde at 4°C overnight and stored in methanol at –20°C. Larvae were then rehydrated to PBST (phosphate buffered saline, 0.1% Tween-20 pH 7.3) in 3 graded steps. After rehydration, embryos were further permeabilized by a 50–55-minute proteinase K treatment (10 μg/mL in PBST), post-fixed for 20 mins with 4% paraformaldehyde at RT, and then washed with PBST (5 times).

### Generation of probes for in situ hybridization of enterocytes

For genes expressed in the intestine and *best4*^+^ enterocytes, antisense *best4*, *otop2*, *pbx3a* and *cdx1b* probes were generated by *in vitro* transcription. Whole-larvae cDNA was generated by isolating total RNA from 3–5 dpf zebrafish larvae using the E.Z.N.A Total RNA kit (Omega, Bio-Tek INC, Cat# R6834-01) and reverse transcribing using the iScript cDNA synthesis kit (BioRad, Cat# 1708891). Fragments of coding sequences of these genes were amplified by PCR using gene-specific primers as follows:

*<u>best4</u>*: 949 bp; F: 5’–TGATGATGGTGGTCTCTGGA and R: 5’–CTTCCAATAGCAGCGTCCAT.
*<u>otop2</u>*: 1011bp; F: 5’–TGATGGCTGTGACTGAGGAG and R: 5’– GTGGTAAACATCGGAATGCC.
*<u>pbx3a</u>*: 958bp; F: 5’–AGCAGGACATCGGAGACATT and R: 5’–AACTGGACGCAGCAGAAGAT.
*<u>cdx1b</u>*: 641bp; F: 5’–CCGTAAGACACCCAAGCCTA and R: 5’– CTCAGCACTACCAGGCAATG.

PCR products were then inserted into the pSC plasmid using the Agilent Strataclone Kit (Cat# 240205) to generate plasmids JFP524 (*best4*), JFP526 (*otop2*), JFP549 (*pbx3a*) and JFP542 (*cdx1b*). Plasmids were linearized with NotI (*pSC-best4*, *pSC-otop2*) or HindIII (*pSC-pbx3a*, *pSC-cdx1b*) and *in vitro* transcribed using T7 (*pSC-pbx3a*, *pSC-cdx1b*) or T3 polymerase (*pSC-best4*, *pSC-otop2*) and fluorescein or digoxygenin RNA labeling kits (digoxygenin: Roche, Cat# 11277073910; fluorescein: Roche, Cat# 11685619910) These reactions were cleaned using the NEB Monarch RNA Cleanup protocol (NEB, Cat# T2040L) and used as anti-sense RNA probes for fluorescent *in situ* hybridization.

### Two color fluorescent *in situ* hybridization

Tyramide-mediated fluorescent *in situ* hybridization was primarily used to characterize *best4*^+^ enterocytes in the zebrafish intestine. Post-fixation, animals were prehybridized in hybridization buffer (50% formamide, 5X SSC, 0.1% Tween-20, 1M citric acid, pH 6.0, 50μg/mL heparin, and 500 μg/mL tRNA) for 2 hours at 70°C in a dry heat block. Hybridization was then performed at 70°C overnight in hybridization buffer with 3 ng/µL of each probe. Following hybridization, the next day, larvae were washed with the following series of buffer washes at 70°C: 75%, 50%, and 25% prehybridization buffer diluted in 2X SSC for 10 minutes each, 2X SSC for 15 minutes, then twice in 0.2X SSC for 30 minutes each. This was followed by a dilution series of 0.2X SSC:PBST (3:1, 1:1, and 1:3 PBST) for 5 minutes each at room temperature. Next, animals in PBST were rocked in 2% blocking buffer (5 g of blocking reagent, Roche, 11096176001) in 1X maleate buffer (150mM maleic acid, 100mM sodium chloride) for at least 2 hours and then incubated in anti-fluorescein-POD F_ab_ fragments (Roche, 11207733910) diluted at 1:400 overnight at 4°C. Upon retrieval, the antibody solution was washed off using PBST and the antibody was developed in dark for 45 minutes without agitation using a tyramide staining solution (TSA PLUS Fluorescein Reagent, Akoya Biosciences, TS-000200) diluted 1:50 in amplification buffer (1X Plus Amplification Diluent, Akoya Biosciences, FP1135). Next, the peroxidase was inactivated in 1% hydrogen for 20 minutes peroxide followed by elution of the antibody with 0.1M glycine (pH 2.2). Larvae were then washed in PBST and incubated in blocking solution for 2 hours at room temperature. The blocking solution was replaced with anti-DIG-POD F_ab_ fragments (Roche 11207733910) diluted 1:500 in blocking reagent. Digoxygenin staining was then developed using the Tyramide PLUS staining solution (TSA PLUS Cy3 reagent, Akoya Biosciences, TS-000202) diluted 1:50 in Amplification buffer for 45 minutes at room temperature. To counterstain DNA, Hoecsht 33342 (Invitrogen, H1399) was added at 1:1000 dilution in PBST and animals were incubated overnight at 4°C then washed 6 times with PBST for 15 minutes each.

### Hybridization Chain Reaction (HCR)

HCR was performed for characterizing pericyte populations, intestinal smooth muscle populations, and the pneumatic duct. HCR probes were designed using the Özpolat lab probe generator (Kuehn et al. 2022), available at https://github.com/rwnull/insitu_probe_generator. Probes were designed with amplifiers B1, B2, B3, and B5, skipping 10 bases from the beginning of the cDNA and by choosing the maximum poly A/T and poly G/C homopolymer length as 5. For each gene, 20 probe pairs were ordered in OPools format (Integrated DNA Technologies) and resuspended in nuclease-free water to a working concentration of 1 µM. Oligos constituting probe sets used in this study are consolidated in Table S3. Following rehydration and fixation, larvae were prehybridized in HCR probe hybridization buffer (Molecular Instruments, Lot# BPH02724) for 0.5–2 hours at 37°C with shaking at 300 rpm. Then, hybridization was performed with 1 µL of each 1 µM probe diluted in 500 µL probe hybridization buffer at 37°C with shaking at 300 rpm overnight (12–16 hours). Probes were washed off using the HCR probe wash buffer (Molecular Instruments, Lot# BPW02624) and subsequently pre-amplified with fresh hairpin amplification buffer (Molecular Instruments) for 30 mins to 2 hours at room temperature. Hairpins (3 µM; Molecular Instruments) were pre-annealed (98°C for 90 seconds and 25°C for 30 mins with a ramp rate of -0.1°C per second) to create hairpin secondary structure. Hairpin mixtures were then diluted 1:100 in hairpin amplification buffer (Molecular Instruments) and larvae were incubated for 12–16 hours at 24°C with 300 rpm shaking. Following amplification, animals were rinsed in 5X SSCT at 24°C followed by washes in PBST and, in some cases, immunostained for a fluorescent protein.

### Immunostaining After HCR

Immunostaining was performed for *flk::mCherry-CAAX* and *flk::GFP* animals to visualize blood vessel architecture in 5 dpf zebrafish larvae. After HCR, samples were further permeabilized with 0.1% Triton X-100 in phosphate buffered saline. Samples were blocked for 2 hours in blocking solution (5% Normal Donkey Serum (Jackson Labs), 10% Bovine Serum Albumin, 1% DMSO, and 0.1% Triton X-100 in phosphate buffered saline). Primary staining was performed with anti-RFP (rabbit, Rockland 600-401-379) and anti-GFP (rabbit, Invitrogen A11122) antibodies diluted 1:1000 in blocking buffer. Secondary staining was performed with goat Alexa Fluor 405-anti-rabbit (Molecular Probes, Cat#: A-31556) diluted 1:1000 in PBST.

### Microscopy and image analysis

Stained larvae were mounted in 1% low melting agarose diluted in 1X Danieau buffer (58 mM NaCl, 0.7 mM KCl, 0.4 mM MgSO_4_, 0.6 mM Ca(NO_3_)_2_, 5 mM HEPES, pH 7.6) and imaged in either a Nikon Eclipse Ti2 inverted microscope with a Nikon DS-R*i*2 camera or an inverted Zeiss LSM 880 Airyscan using a 10X air, 20X air, and 40X long working distance water/oil objective. Image processing was performed in ImageJ/Fiji and Photoshop (Adobe). The acquired z-stacks were projected using Fiji, and brightness and contrast were set in Fiji using only a linear relationship in the lookup table. Cropping and resizing was performed using Photoshop (Adobe) and figures were assembled using Illustrator (Adobe).

### Inference of developmental trajectories using URD

Transcriptional trajectories were constructed using URD v1.1.2 as previously described (Farrell et al. 2018) from the putatively non-neural crest derived (i.e. *foxc1a/foxc1b* negative and *prrx1a/b* negative) viSMCs, vaSMCs, and myofiboblasts and intestinal derivatives to determine the molecular events that occur as cells diversify and differentiate in these tissues. Cells belonging to each group were selected by cluster identities and used to create a URD object using the URD::seuratToURD2() function. Previously identified highly variable genes were maintained.

To identify and remove outlier cells, a *k*-nearest neighbor network was calculated between cells using Euclidean distance in gene expression space with 100 nearest neighbors. Cells were then removed based on their distance to their nearest neighbor or unusually high distances to the 20^th^ nearest neighbor using the function URD::knnOutliers (mural-cells: x.max = 25, slope.r = 1.1, int.r = 4, slope.b = 1.85, int.b = 8; intestine: x.max = 23, slope.r = 1.2, int.r = 5, slope.b = 0.85, int.b = 7.5).

URD uses user-defined starting (’root’) and endpoints (‘tips’) for building trajectories. The earliest stage cells from each subset (36–58 hpf for SMCs/pericytes and 14–21 hpf for intestinal subtypes) were selected as the ‘root’ or starting point of the tree. Endpoints were defined from late stage cells (120 hpf from the intestinal tissues and 108–120 hpf for the SMCs/pericytes), based on their cluster identity. Clusters were excluded that represented clear progenitor or precursors based on gene expression of cell cycle genes (e.g., intestine: cluster 14).

To identify cells along trajectories between the ‘root’ and each ‘tip’, first a diffusion map was calculated using the function URD::calcDM with parameters nn = 100, sigma = 12 (non-skeletal muscle) or sigma = 8.2 (intestine), which draws on the R package *destiny* (Haghverdi, Buettner, and Theis 2015). Next, pseudotime was computed using the function URD::floodPseudotime (n = 100, minimum.cells.flooded = 2). Then, biased random walks were simulated starting from each terminal using the function URD::simulateRandomWalk, with the following parameters—SMCs/pericytes: optimal cells forward = 10, max.cells.back = 20, n.per.tip = 25000, root.visits = 1, max.steps = 5000; intestine: optimal cells forward = 20, max.cells.back = 40.

Next, we fit a branching tree structure to each set of trajectories using the URD::buildTree command with the following parameters—SMCs/pericytes: divergence.method = “preference”, cells.per.pseudotime.bin = 10, bins.per.pseudotime.window = 8, p.thresh = 0.001; intestine: divergence.method = “preference”, cells.per.pseudotime.bin = 25, bins.per.pseudotime.window = 5, p.thresh = 0.1. To visualize the trajectories, we generated force-directed layouts on cells that had been robustly visited during the random walks (i.e., cells that have a minimum visitation frequency of 0.5) using the command URD::treeForceDirectedLayout function with the following parameters num.nn = 87 (intestine) or num.nn = 110 (SMCs/pericytes), cells.to.do = robustly.visited.cells, cut.outlier.cells = NULL, cut.outlier.edges = NULL, cut.unconnected.segments = 2, min.final.neighbors = 4.

### Identifying gene cascades along developmental trajectories

Gene cascades were constructed using two different methods for intestinal SMCs and intestinal derivatives. For intestinal SMCs, each major group (or clade) of branches from the end of the branching tree were considered as a single entity and compared against each other pairwise to identify differentially expressed genes. In addition, using URD’s branching tree as a framework, the URD::aucprTestAlongTree() function was also used to find genes that are differential markers of each lineage within each group using the parameters: must.beat.sibs = 0.6, auc.factor = 1.1, log.effect.size = 0.4. These differentially expressed genes were further curated based on other criteria as described previously (Raj et al. 2020). All genes that had a fold change of 0.6 along the trajectory pursued and a classifier score of 1.05 were used for downstream analysis. Genes along the individual intestinal SMCs branches were normalized to the maximum observed for each gene within this tissue and ordered based on the pseudotime that they enter “peak” expression (defined as 50% higher expression than the minimum expression value). Gene expression dynamics within the intestinal SMCs were fit using smoothed spline curves using the function URD::geneSmoothFit() with parameters method = “spline”, spar = 0.5, moving.window = 5, cells.per.window = 8, pseudotime.per.window = 0.005. For the intestinal SMC trajectory, genes were further compared between the two branches to select genes that are specific to one branch or another or markers of both (i.e., this strategy was used to differentiate between intestinal SMC precursors, circular and longitudinal SMC markers).

In the intestine trajectory, for each intestinal population, cells in each segment were compared pairwise with cells from each of the segment’s siblings and children using the function URD::aucprTestAlongTree(log.effect.size = 0.4, auc.factor = 0.6, max.auc.threshold = 0.85, frac.must.express = 0.1, frac.min.diff = 0, must.beat.sibs = 0.6). Genes were called differentially expressed if they were expressed in at least 10% of cells in the branch under consideration and were 0.6 times better than a random classifier for the population as determined by the area under a precision-recall curve. Gene expression of each marker gene in each trajectory within the intestine was then fit using an impulse model using the function URD::geneCascadeProcess() with parameters: moving.window = 5, cells.per.window = 18, pseudotime.per.window = 0.01. Cells in each trajectory were grouped using a moving window through pseudotime within which mean expression was calculated. Expression was then normalized to the maximum observed expression for each gene within the intestine. The parameters of the impulse model were then used to calculate the onset time for each gene and order genes according to the pseudotime of their expression onset.

### Simulation of artificial pericyte doublets

To simulate artificial pericyte doublets, we mixed gene expression signatures of the general pericyte cluster (pericyte-0: C9) that did not express any unique markers with that of other cell types that expressed the characteristic markers of the two pericyte populations identified in our study (C20: pericyte-1; C4: pericyte-2). We identified the top 3 genes expressed by C20 and C4 respectively and extracted cells from the global dataset other than pericytes that strongly expressed these genes (log-normalized expression >2). Out of these cells, only ones between 60–120 hpf were used for downstream analysis as the two putative pericyte clusters consisted of cells encompassing those stages. For each group, we then randomized the pool of C9 (pericyte-0) cells and the extracted non-pericyte cells to create 5000 random cell pairs to use for creating artificial doublets. To create a mixed gene expression signature characteristic of doublets in scRNAseq, for each cell pair, we unlogged the expression data, took the mean of the pericyte-0 and non-pericyte cells’ expression, and re-calculated the log. We limited the genes used to calculate average expression to the union of the highly variable genes calculated on the global dataset and the non-skeletal muscle atlas. To understand how similar or different these doublets are to the distinct pericyte subpopulations, we calculated Euclidean distances in variable gene expression space for cells within a pericyte population (i.e. pericyte-1 to pericyte-1) and between a pericyte population and its simulated doublets (i.e. pericyte-1 to pericyte-1-simulated-doublets) using the “dist” function. Simulated doublets were always more distant from the pericyte subpopulations than other cells within the population.

To further confirm that the pericyte-1 and -2 clusters are not artefacts, we further performed differential gene expression analysis between the individual pericyte clusters versus their corresponding artificial doublet signatures. For each pericyte cluster, based on the distances calculated above, we chose the nearest 5% artificial doublet cells (*i.e.* those most similar to the putative pericyte population) and compared their gene expression using the Seurat::FindMarkers function. Even the most similar simulated doublets did not recapitulate the expression of the pericyte-1 and pericyte-2 populations.

### Comparison of gene expression between species

Gene expression comparisons were performed against a human colon (Parikh et al. 2019) and a small intestine (Burclaff et al. 2022) scRNAseq dataset. Gene orthologs between zebrafish and human were obtained using the R Bioconductor package “biomaRt”. Genes for which orthologs were not present in one of the two datasets (e.g., human *GUCA2A*, *GUCA2B*, zebrafish *sctr*, *tacr2*) were also included in the cross-species analysis.

## Supporting information

Supplementary Table 1 (Cluster Annotations)

Supplementary Table 2 (Gene Module Annotations)

Supplementary Table 3 (HCR Probe Sequences)

## Acknowledgements

Funding for this study was provided by the NICHD Intramural Program to JAF (ZIAHD008997) and the Allen Discovery Center for Lineage Tracing and the NIH to Alexander Schier (R01HD085905, DP1HD094764). We thank Alexander Schier for his generosity in supporting early aspects of this project and his excellent mentorship. We thank Chris McGinnis the Gartner lab for gifting MULTI-seq reagents. We thank Dan Castranova and Brant Weinstein for assistance in the proper identification and imaging of perivascular cells and vessels. We thank stellar core facilities and personnel: the Harvard Bauer Core Facility, NICHD Molecular Genomics Core, NICHD High Performance Computing and Biowulf team, NIH Shared Zebrafish Facility, Ryan Dale, Nikki Swan, Loc Vu, and Matt Breymaier. We thank Jamie Gagnon, Katherine Rogers, Brant Weinstein, Harry Burgess, and members of the Farrell and Rogers labs for helpful comments on the manuscript.

## Author Contributions

JAF, YW, and AS conceived the study. AS and JAF wrote the manuscript. AS, GM, and JF performed the data analysis. AS and JF annotated single-cell data. JAF and YW collected single-cell transcriptomes. PC and AS performed other experiments, including RNA *in situ* hybridizations. JAF built Daniocell.

## Data and materials availability

Daniocell is accessible at http://daniocell.nichd.nih.gov/. Sequencing data is available as FASTQs and UMI count tables under NCBI GEO accession GSE223922. Code will be available at https://github.com/ farrelllab/2023_Sur/.

