## Supplementary Figures and Legends for "Single-cell analysis of shared signatures and transcriptional diversity during zebrafish development"

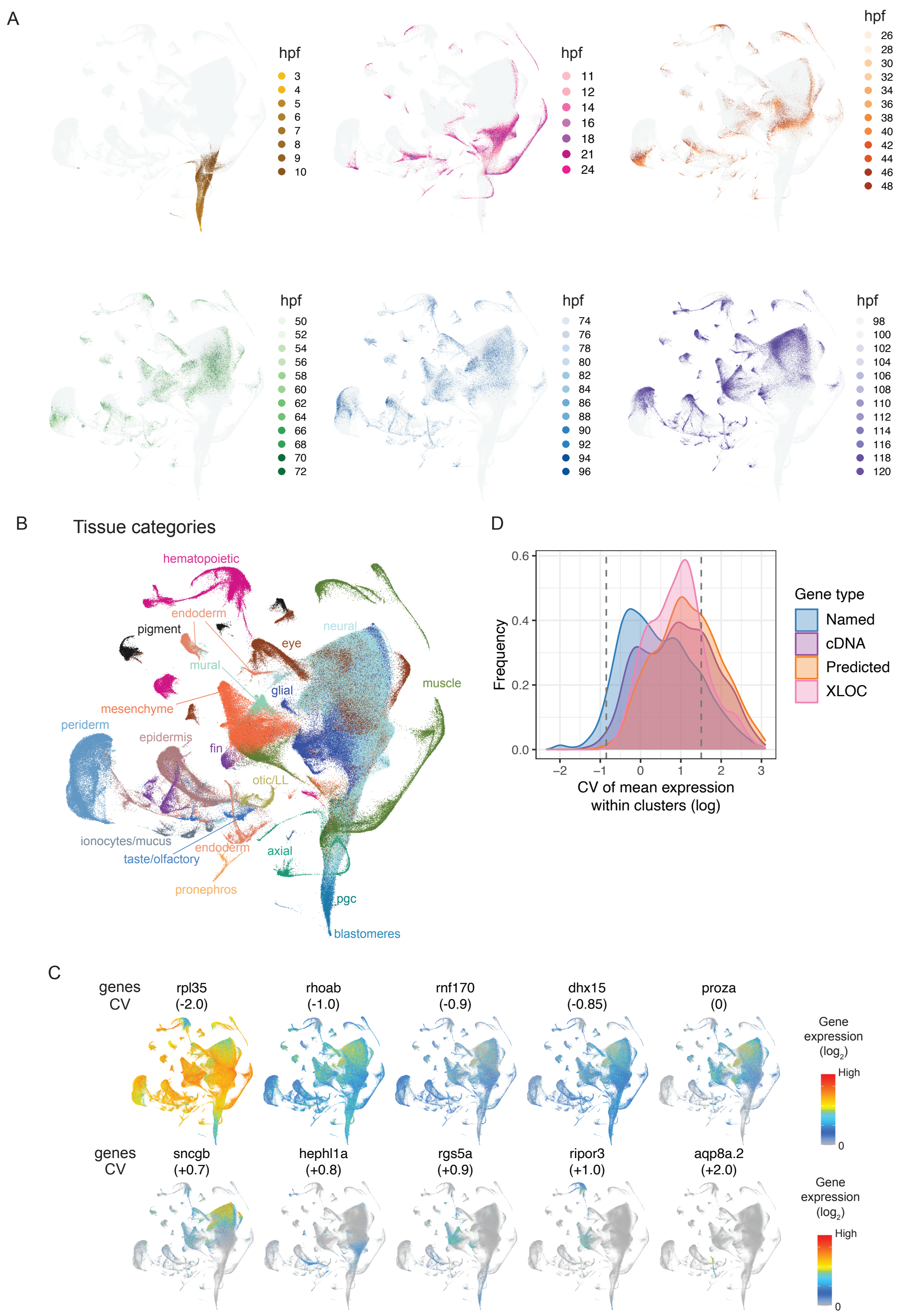


**Supplementary Figure 1. Categorization of cells into distinct tissue types and understanding gene expression variation. (A)** UMAP projection of single-cell transcriptomes, colored by developmental stage (colored as in Fig. 1A–B) but shown as groups of stages across multiple plots. **(B)** Single cells colored according to the 19 different tissue categories that were used for downstream analysis. **(C)** Example expression patterns for genes with different log CV values (as plotted in Fig. 1d) demonstrate how animal-wide expression patterns are represented using this metric. **(D)** Distribution of the coefficient of variation of cluster means of expression for each gene (log-transformed) for genes of different classes: (1) 'named' genes, (2) unnamed cDNA clones (*e.g. si:dkey-, si:ch211-, zgc:*, and others), (3) computationally predicted genes from Refseq (LOC) and Ensembl (e.g. *BX-, CABZ-*, and several others), and (4) transcripts introduced in the Lawson annotation from a *de novo* transcriptome assembly (XLOC-).

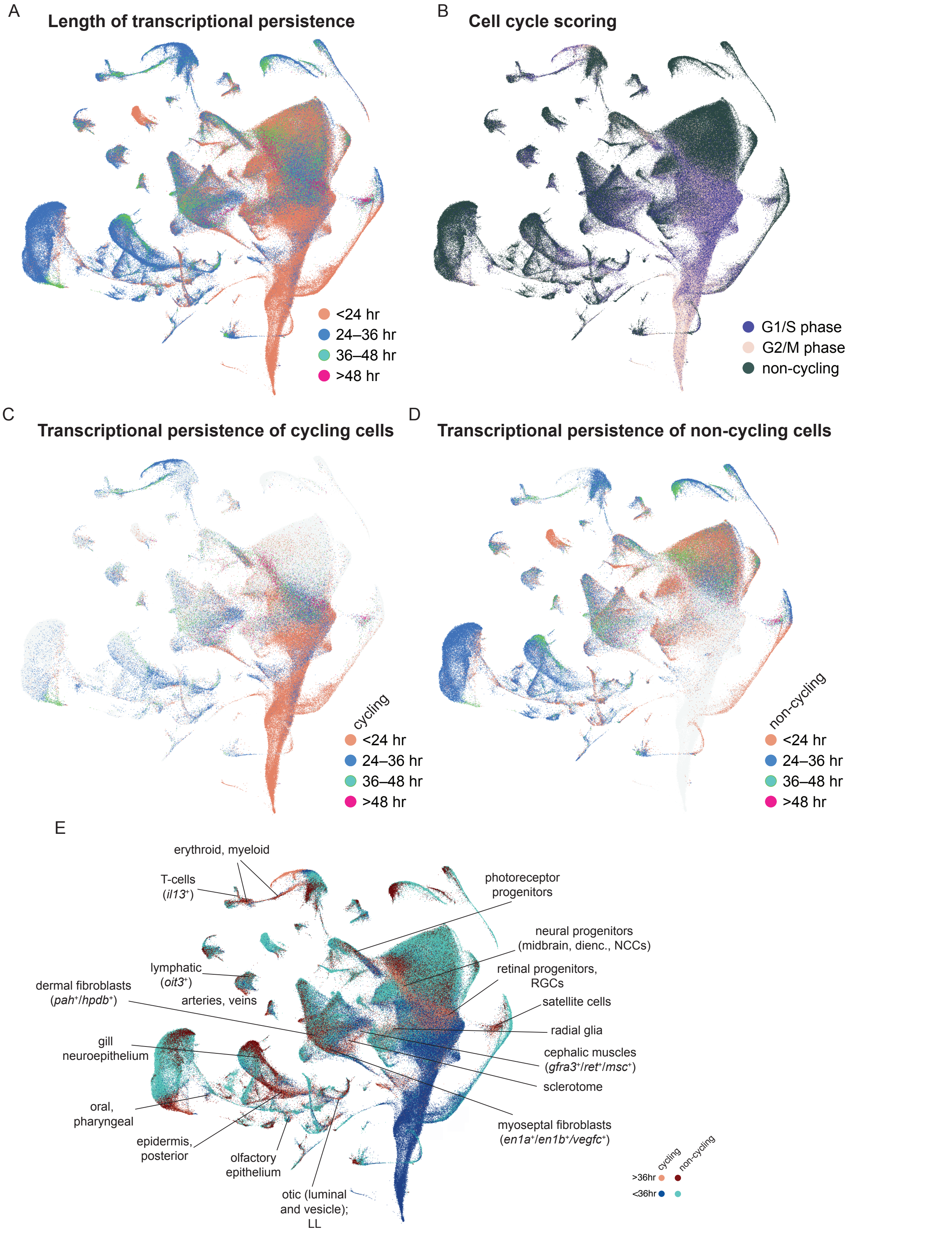


**Supplementary Figure 2**: **Persistence of transcriptional states during zebrafish development. (A)** For each cell, as a measure of the duration of its transcriptional state, shown is the range of developmental time occupied by its ε-nearest neighbors. **(B)** Cell-cycle state of all cells, as determined by transcription of cell cycle-related genes. **(C, D)** Similar to panel A, duration of cells' transcriptional state separately for (**C**) cycling or (**D**) non-cycling cells. **(E)** UMAP projection showing categorization of cycling and non-cycling cells based on whether they exhibit a stable transcriptome for ≥36 hours or ≤36 hours. Populations of proliferating cells whose transcriptional state persists ≥36 hours (also called "long-term" transcriptional states) are labeled on the plot.

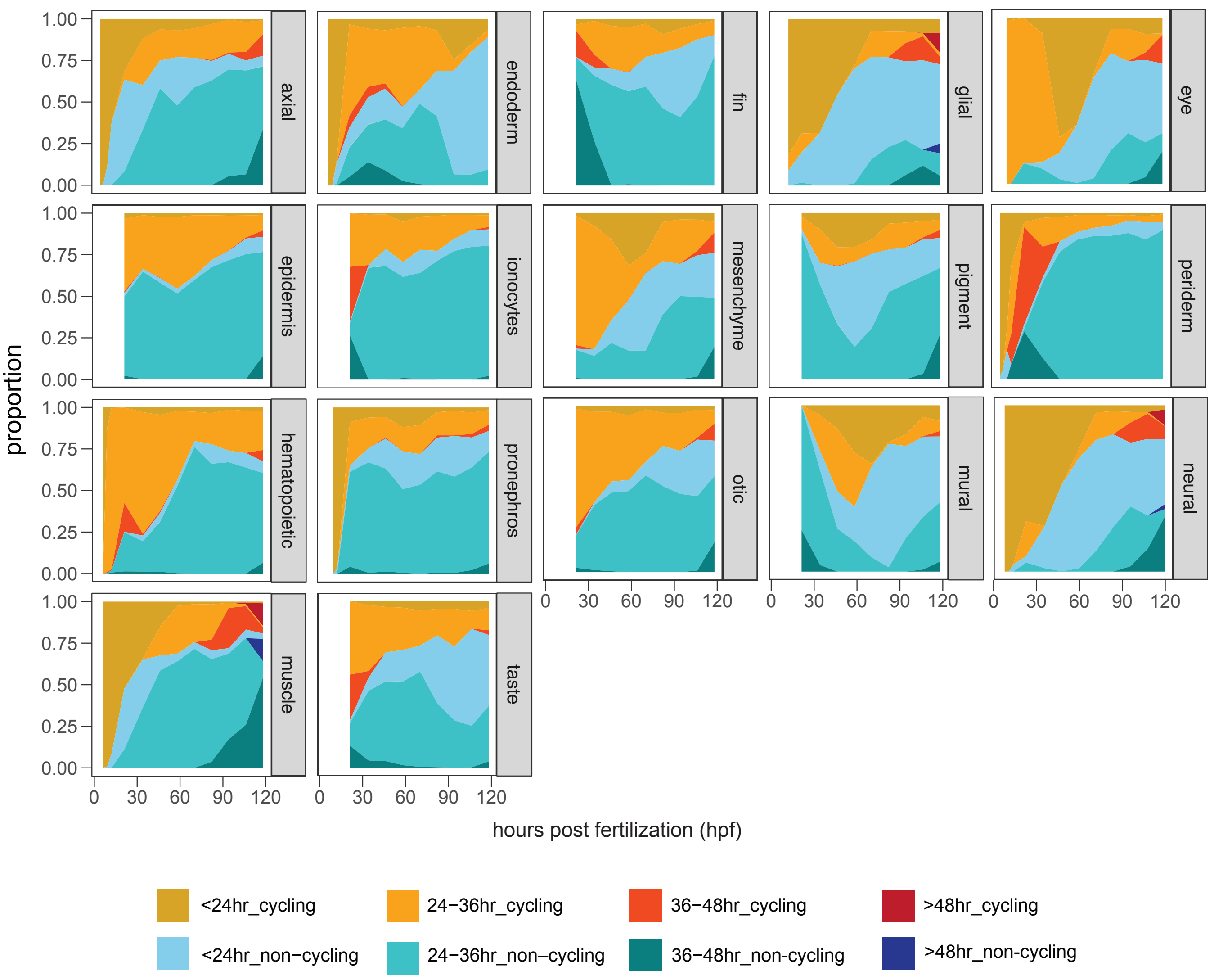


**Supplementary Figure 3**: **Persistence of transcriptional states during development within individual tissues.** Duration of cells' transcriptional states (as shown in Fig 1E–F, Supplementary Figure 2), based on the range of developmental time occupied by their ε-nearest neighbors. Cells are grouped by the tissue categories that are displayed in Supplementary Figure 1B and used for iterative clustering analysis. Categories are based on cells' cell cycle status ("cycling" or "non-cycling") and transcriptional persistence. Blastomere and PGC categories were not analyzed in this manner due to either limited developmental duration or limited cell count.

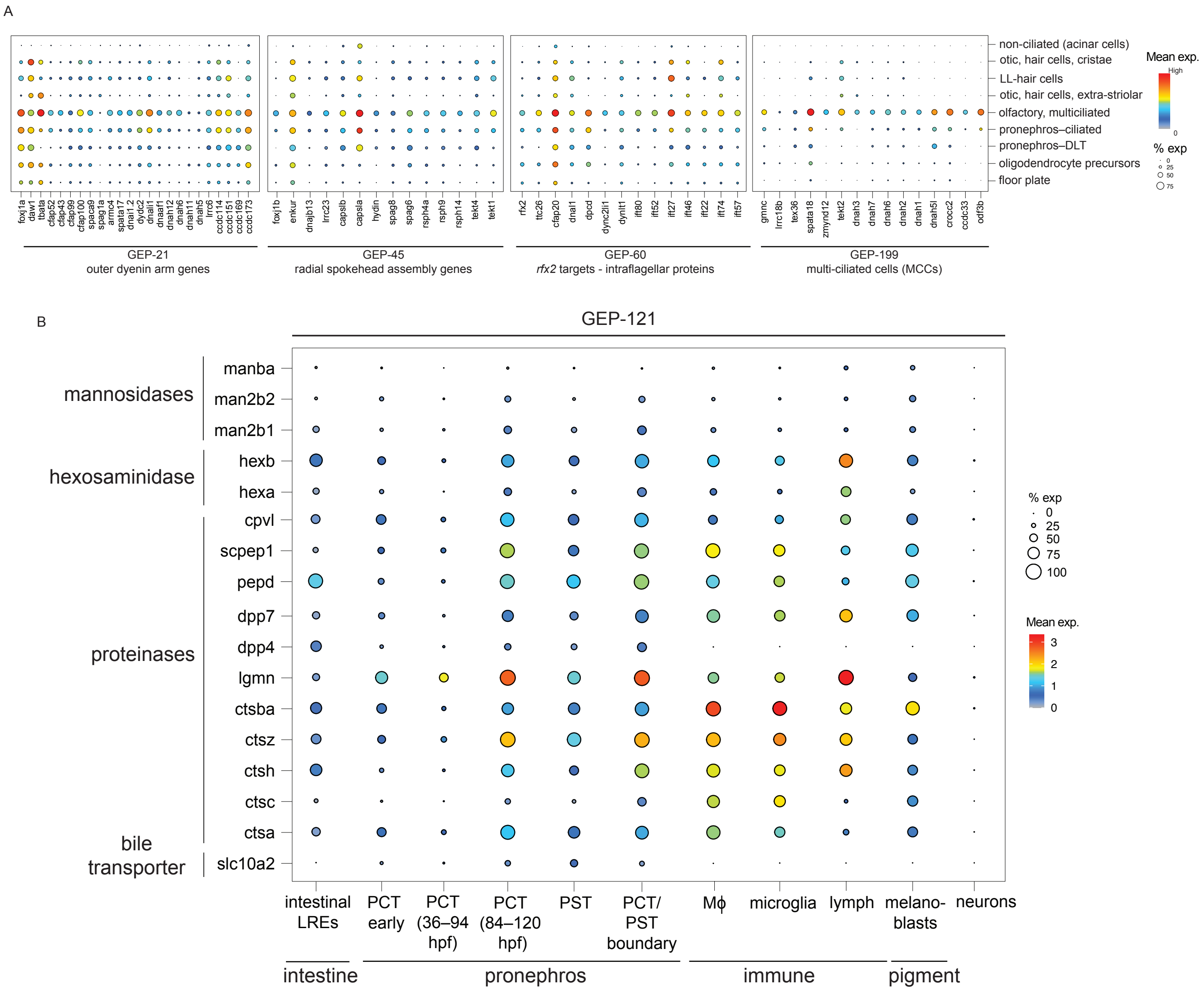


**Supplementary Figure 4**: **Ciliary gene modules and a lysosomal degradation module shared between lysosome-rich cells. (A)** Dot plot showing genes in four ciliary gene modules, annotated with putative functions. X-axis: gene members of modules GEP-21, GEP-45, GEP-60, and GEP-199. Y-axis: ciliated cell types, and acinar cells as a non-ciliated control. **(B)** Dot plot showing lysosomal degradation gene members of GEP-121. X-axis: cell clusters, including pronephros proximal tubule segments, lysosomal-rich enterocytes, macrophages, pigment-cells, and neurons as a non-lysosome-rich control. **LREs** – lysosome-rich enterocytes**; PCT** – proximal convoluted tubule; **PST** – proximal straight tubule; **Mϕ** – macrophages; **hpf** – hours post-fertilization; **GEP** – gene expression program.

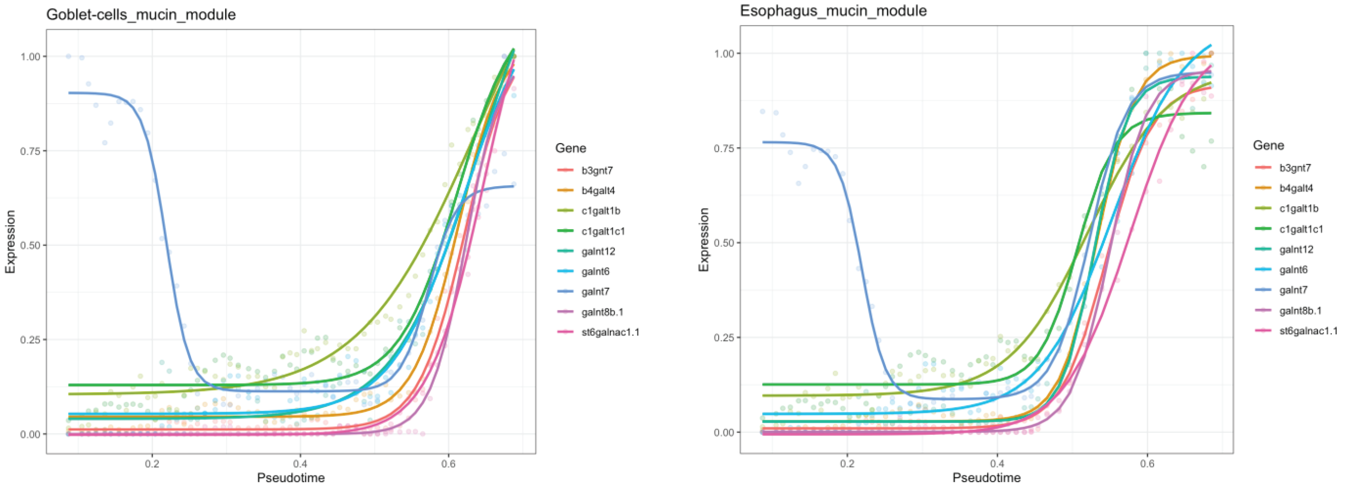


**Supplementary Figure 5**: **Mucin O-glycosylation genes exhibit similar expression kinetics in multiple gut cell types**. Expression kinetics of mucin module (GEP-94) genes within either the intestinal goblet cells or esophageal mucous-secreting cells, fit with an impulse function. X-axis: pseudotime calculated within the gut (a measure of developmental progression); Y-axis: Gene expression, scaled to maximum expression within each tissue.

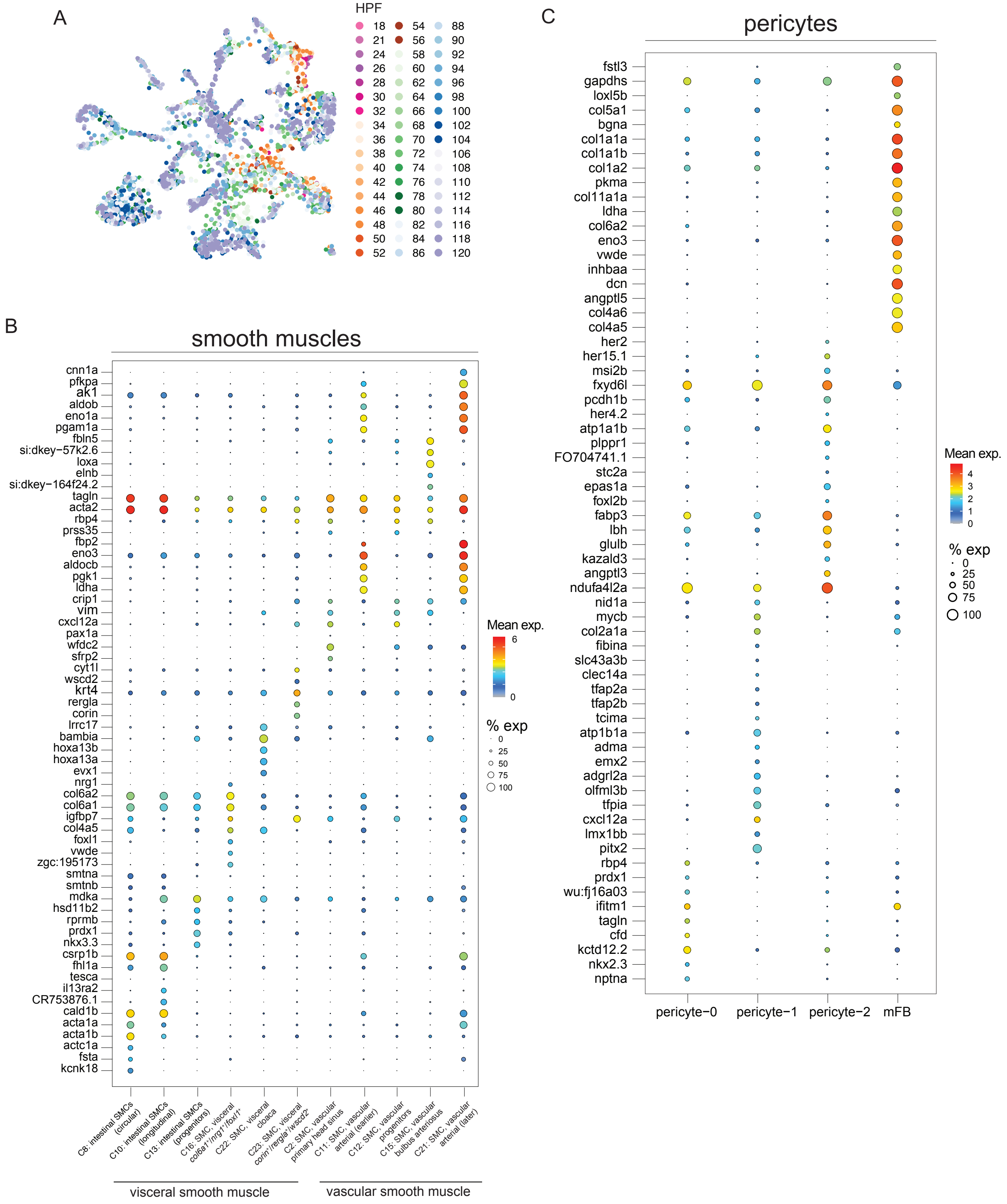


**Supplementary Figure 6: Differential gene expression within smooth muscle and pericyte cell populations. (A)** UMAP projection of 3,866 non-skeletal muscle cells, color-coded by developmental stage. **(B)** Dot plot showing top differentially expressed genes between smooth muscle populations (including visceral and vascular) captured in our dataset. **(C)** Dot plot showing top differentially expressed genes between the pericyte subpopulations in the zebrafish head. Note that "pericyte-0" cluster does not express any unique genes indicating a general pericyte population. **mFB** – myofibroblasts; **SMC** – smooth muscle cell; **HPF** – hours post-fertilization.

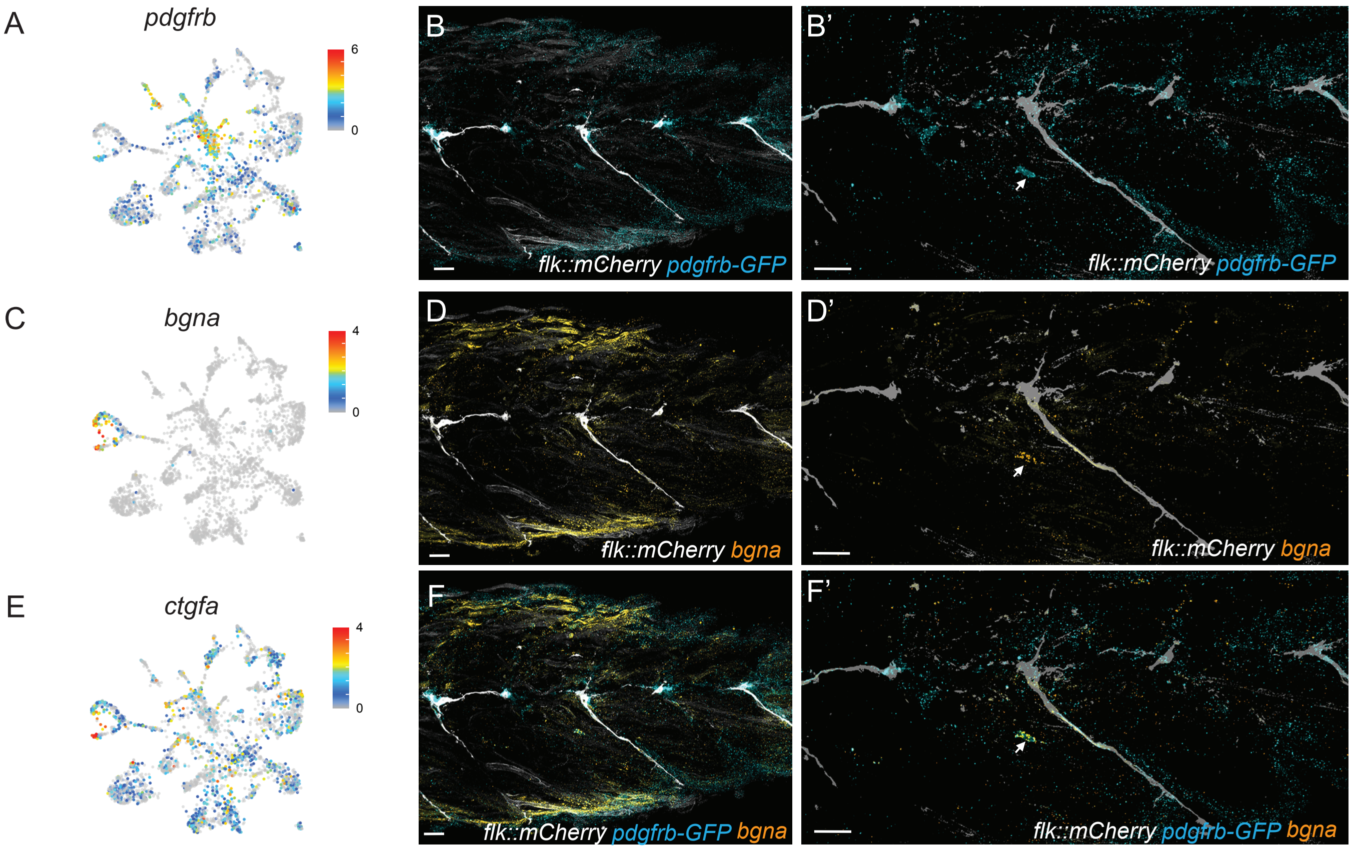


**Supplementary Figure 7**: **Characterization of a myofibroblast (C3: myofibroblasts) population near zebrafish trunk vasculature. (A, C, E)** Expression of genes (*pdgfrb*, *bgna*, and *ctgfa*) expressed in the myofibroblast cluster (C3) visualized on UMAP projections. Color bar represents mean gene expression levels for each gene **(B, D, F)** RNA *in situ* hybridization for markers specific to the myofibroblast cluster (*bgna*) along with a general perivascular marker (labeled by *pdgfrb::GFP)* and vasculature (labeled by *flk::mCherry-CAAX)* in 5 dpf larval trunk. Expression of *bgna* was observed in *pdgfrb::GFP*+ cells within trunk muscle. Scale bar: 25 µm.

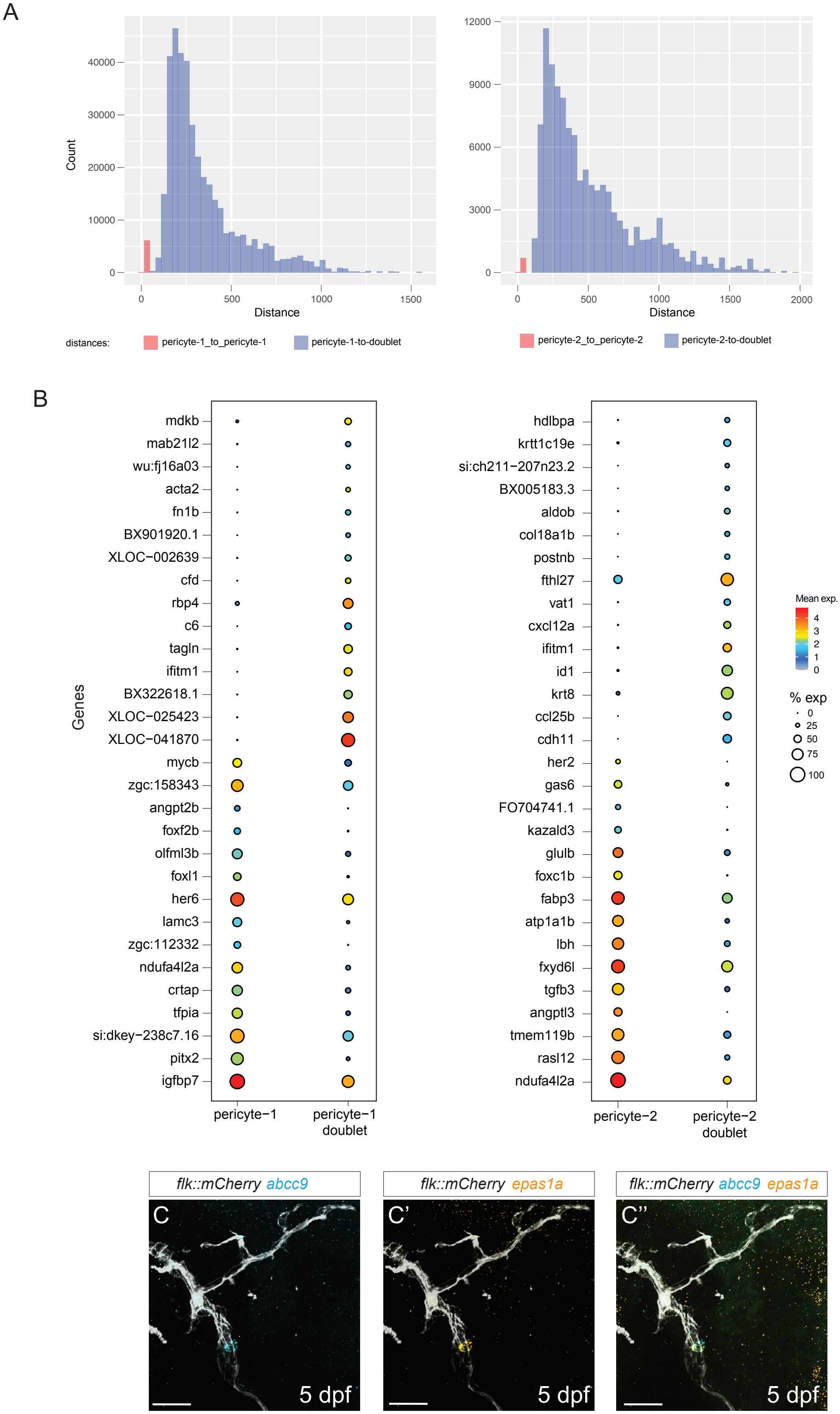


**Supplementary Figure 8: Simulation of artificial pericyte doublets reveals distinct gene expression signatures of the two pericyte populations. (A)** Histogram showing the comparative distribution of Euclidean distances in variable gene expression space between each pericyte population (C20 and C4) to themselves (i.e., distances between C20 cells to C20 cells, and distances between C4 cells to C4 cells) compared to distances between each pericyte population and artificial doublets created between pericyte-0 cells and other populations that express top characteristic markers of C20 and C4 respectively (see Methods). Colors represent the two categories of Euclidean distances computed. For each pericyte population, no artificial doublets were perfectly similar to these populations, suggesting they do not reflect incompletely dissociated cells. **(B)** Dotplot showing differentially expressed markers between the artificial doublets and each pericyte cluster indicating that pericyte-1 (C20) and pericyte-2 (C4), demonstrating that the artificial doublets do not recapitulate the expression of several genes that characterize the pericyte transcriptional states. **(C–C")** RNA *in situ* hybridization for markers specific to the pericyte-2 population (*epas1a*) on a *flk::mCherry-CAAX* background in 5 dpf larvae co-stained with a characteristic pericyte marker *abcc9*. A magnified image of the posterior cerebral vein similar to Fig. 3F–G" shown. Scale bar: 25 µm.

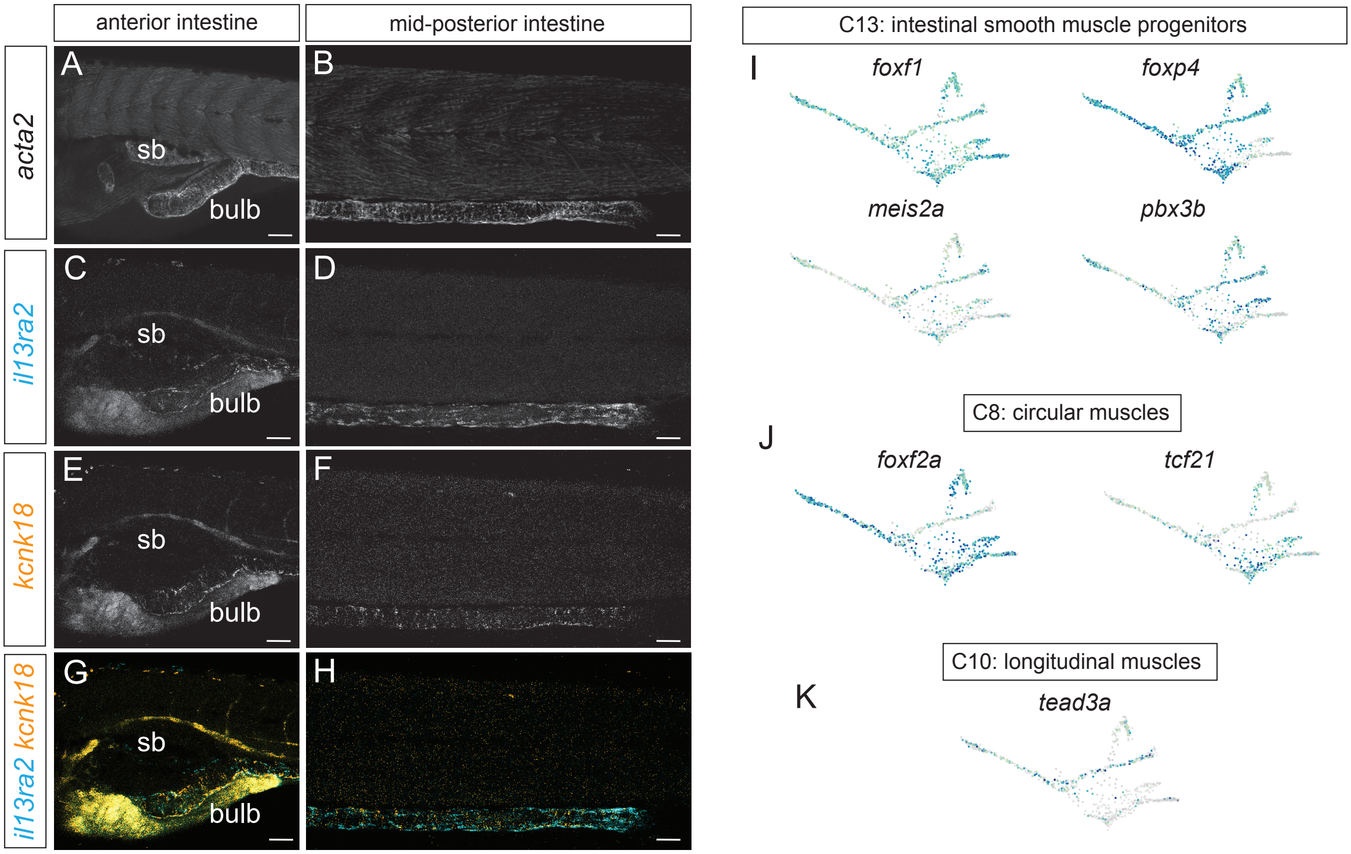


**Supplementary Figure 9**: **Distinct markers label the circular and longitudinal SMCs lining the zebrafish intestine. (A–H)** RNA *in situ* hybridization of markers expressed in longitudinal (*il13ra2*) and/or circular smooth muscle cells (*kcnk18*) lining the zebrafish intestinal tract. (A, C, E, G) anterior intestine view, including the intestinal bulb. (B, D, F, H) mid- to posterior intestine view. (I–K) Expression of intestinal smooth muscle layer-specific TFs for the common progenitors (C13), circular (C8), and longitudinal (C10) visualized on the URD trajectory. Scale bar: 50 µm

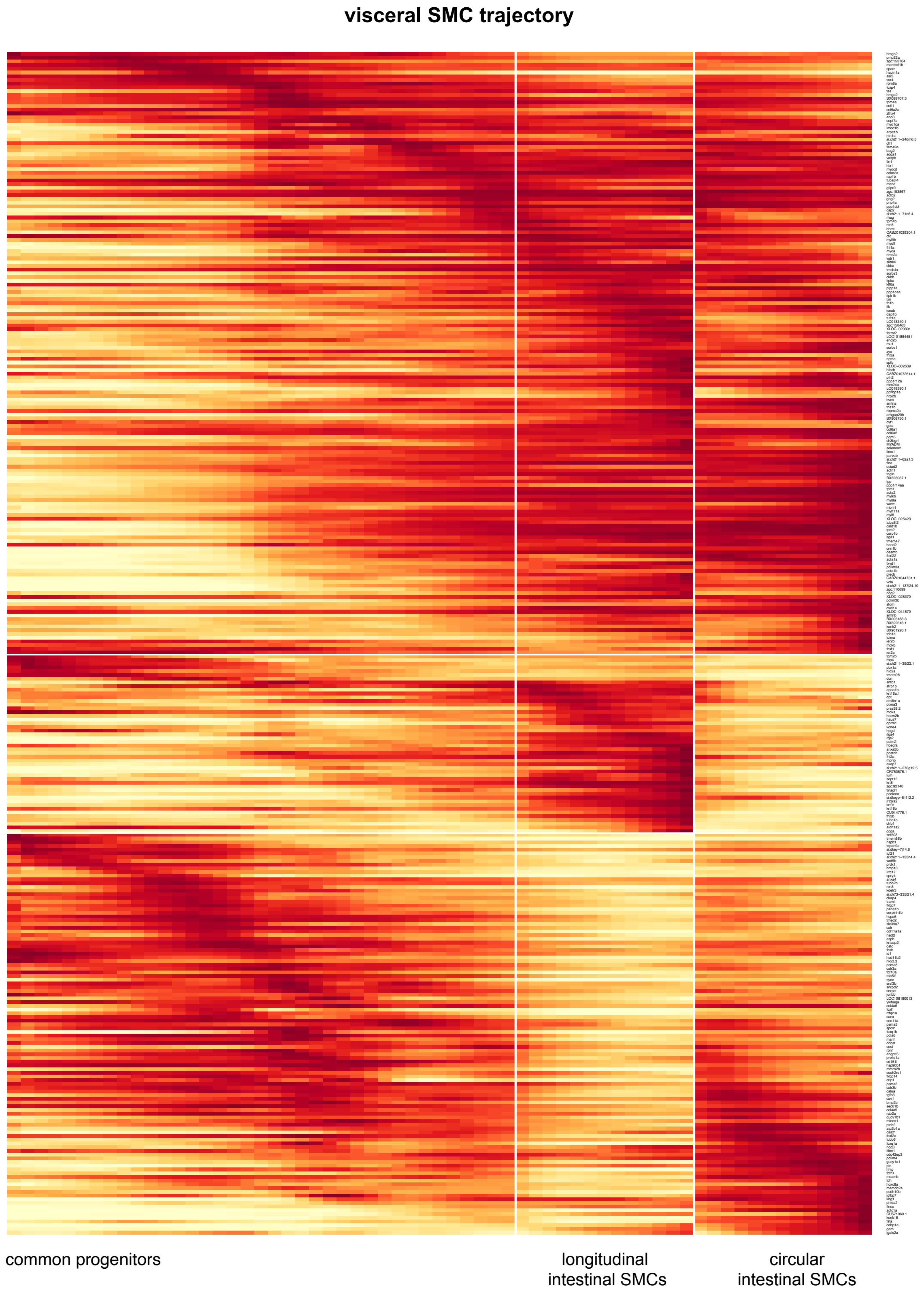


**Supplementary Figure 10**: **Gene expression dynamics during viSMC development**. Heatmap of expression dynamics of genes that vary significantly along URD trajectories to intestinal longitudinal and circular SMCs. X-axis: Pseudotime. Left section represents viSMC progenitors upstream of branchpoint; two right sections are viSMC cells after branchpoint between longitudinal and circular smooth muscle. Color: yellow is low expression, red is high expression (scaled to maximum observed expression).

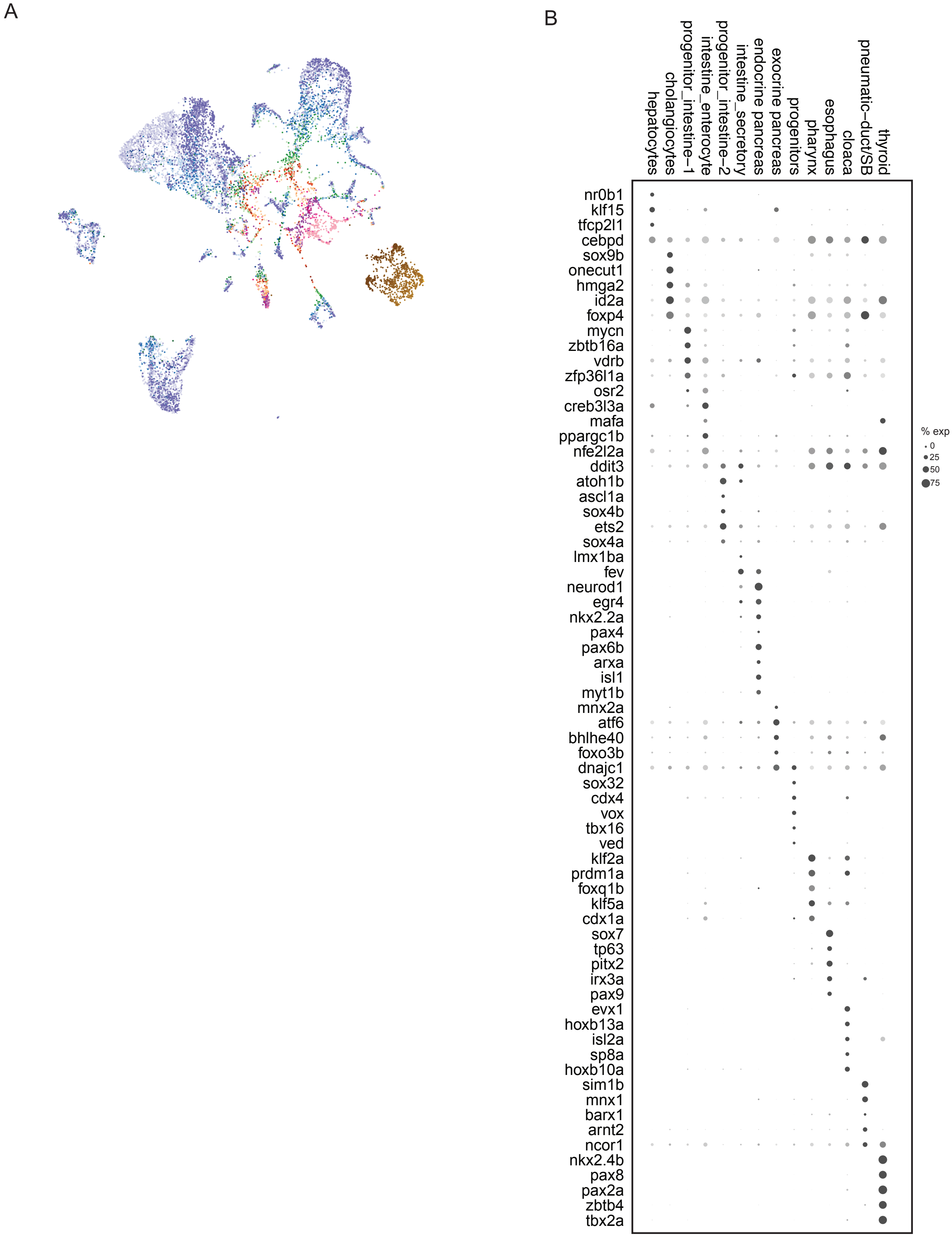


**Supplementary Figure 11**: **Distinct transcriptional profiles of endodermal derivatives.** (**A**) UMAP projection of 12,592 endodermal cells during zebrafish development, colored by developmental stage. Colors corresponding to developmental stages is similar to that shown in Fig. 1a. (**B**) Dot plot of differentially expressed transcription factors across endodermal derivatives. Color: mean expression per cell type (black – high expression); size: percent of cluster cells that express gene, x-axis: endodermal cell types each transcription factor is expressed in.

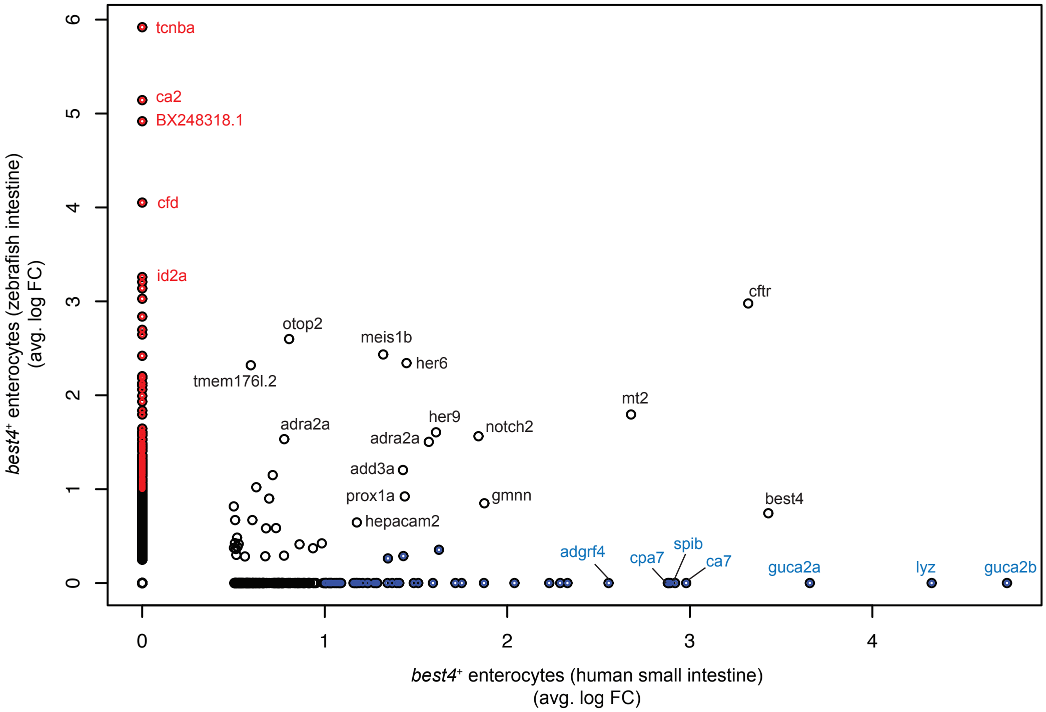


**Supplementary Figure 12**: **Shared gene expression between zebrafish and human small intestinal *best4*+ enterocytes**. Similar to Fig. 6B. Average log-fold enrichment of genes in *best4*+ enterocytes compared to *best4*+enterocyte subtypes in human small intestine (x-axis, Burclaff et al. 2022) and zebrafish intestine (y-axis). Blue: human small intestine-specific *best4*+ enterocyte markers; red: zebrafish-specific *best4*+ enterocyte markers; black: shared markers.

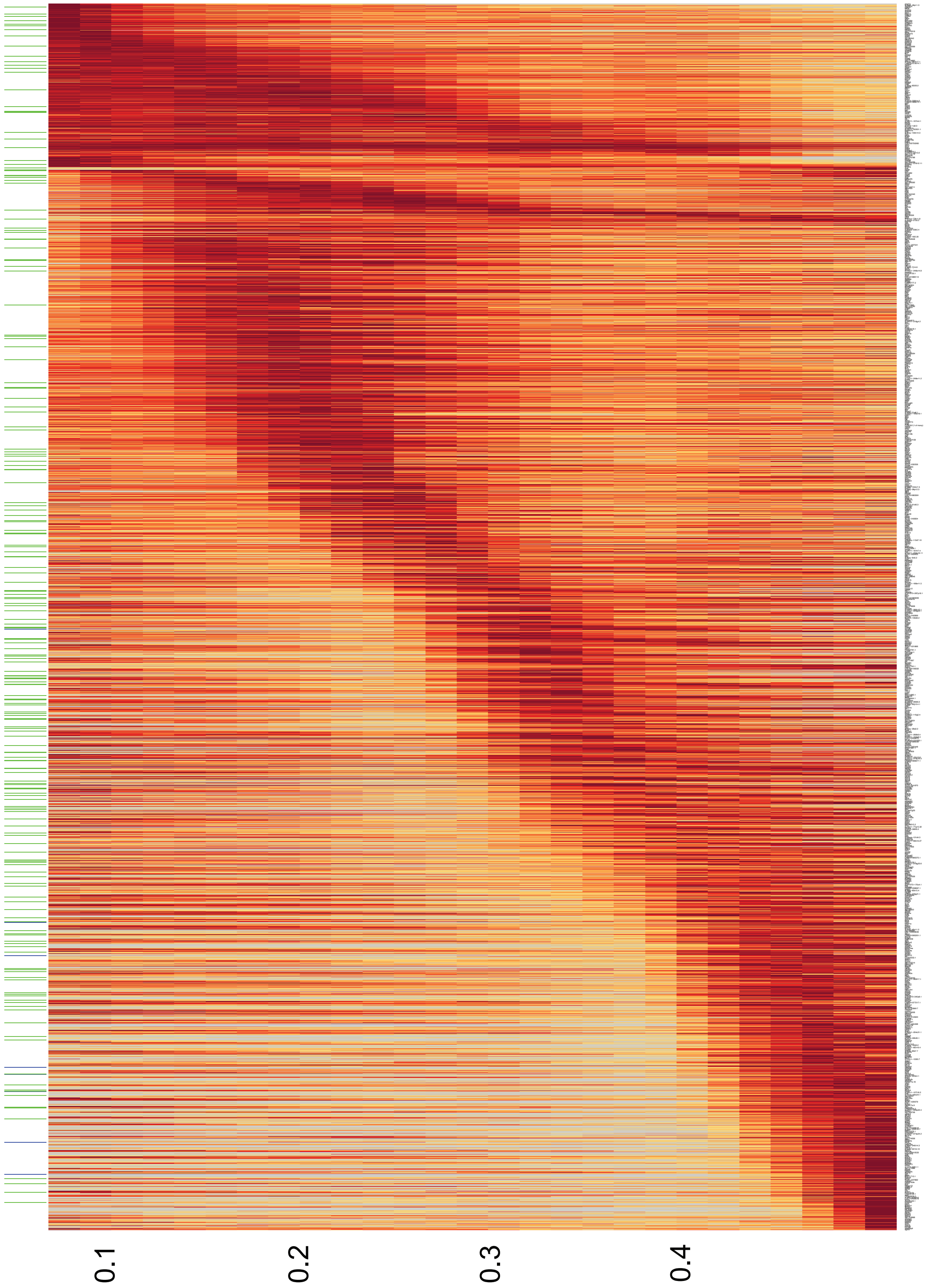


**Supplementary Fig. 13**: **Gene expression during *best4*+ enterocyte specification**. Heatmap of gene expression dynamics along trajectory to *best4*+ enterocytes. X-axis: pseudotime. Green bars on the left indicate transcription factors that are expressed along the cascade to *best4*+ cells. Blue bars indicate *best4*+ enterocyte specific markers expressed around the end of the cascade.
